# Tumor cell heterogeneity drives spatial organization of the intratumoral immune response in squamous cell skin carcinoma

**DOI:** 10.1101/2023.04.25.538140

**Authors:** Miho Tanaka, Lotus Lum, Kenneth Hu, Cecilia Ledezma-Soto, Bushra Samad, Daphne Superville, Kenneth Ng, Zoe Adams, Kelly Kersten, Lawrence Fong, Alexis J. Combes, Matthew Krummel, Melissa Reeves

## Abstract

Intratumoral heterogeneity (ITH)—defined as genetic and cellular diversity within a tumor—is linked to failure of immunotherapy and an inferior anti-tumor immune response. The underlying mechanism of this association is unknown. To address this question, we modeled heterogeneous tumors comprised of a pro-inflammatory (“hot”) and an immunosuppressive (“cold”) tumor population, labeled with YFP and RFP tags respectively to enable precise spatial tracking. The resulting mixed-population tumors exhibited distinct regions comprised of YFP^+^ (hot) cells, RFP^+^ (cold) cells, or a mixture. We found that tumor regions occupied by hot tumor cells (YFP^+^) harbored more total T cells and a higher frequency of Th1 cells and IFN*γ*^+^ CD8 T cells compared to regions occupied by cold tumor cells (RFP^+^), whereas immunosuppressive macrophages showed the opposite spatial pattern. We identified the chemokine CX3CL1, produced at higher levels by our cold tumors, as a mediator of intratumoral macrophage accumulation, particularly immunosuppressive CD206^Hi^ macrophages. Furthermore, we examined the response of heterogeneous tumors to a therapeutic combination of PD-1 blockade and CD40 agonist on a region-by-region basis. While the combination successfully increases Th1 abundance in “cold” tumor regions, it fails to bring overall T cell activity to the same level as seen in “hot” regions. The presence of the “cold” cells thus ultimately leads to a failure of the therapy to induce tumor rejection. Collectively, our results demonstrate that the organization of heterogeneous tumor cells has a profound impact on directing the spatial organization and function of tumor-infiltrating immune cells as well as on responses to immunotherapy.

## Introduction

Immunotherapies have shown great promise in treating cancer by activating a patient’s own immune cells to fight their tumor, but only a fraction of patients respond to current treatment regimens^1–4^. Immune checkpoint blockade (ICB) therapy routinely achieves durable cures in only a limited number of cancer types, and in many tumor types, response rates are as low as 15% with high relapse rates^3, 4^. Genetic heterogeneity within a tumor is a significant factor that has been linked to poor ICB response^5–7^. Tumors are often comprised of multiple populations of cancer cells, with each population carrying a distinct set of genetic alterations and phenotypic behaviors^8–10^. Genetic heterogeneity—often given the generalized label “intratumoral heterogeneity” (ITH)—is inherent in all cancer types^6^, and studies in multiple cancer types have shown that tumors with high genetic heterogeneity (high ITH) are less likely to respond to ICB regimens^5, 7^. Further, in patients, tumors are frequently a patchwork of “hot” and “cold” regions^11–13^—characterized by high and low T cell infiltrates, respectively —indicating that a dysfunctional T cell phenotype can be spatially localized^14^. One study of 85 pre-treatment lung cancers revealed that over two-thirds of tumors investigated contained both hot and cold regions^11^. However, model systems that lend themselves to mechanistically interrogating the impact of tumor heterogeneity on the spatial organization of immune cells within a tumor are limited at present.

Heterogeneity-induced impairment of the anti-tumor immune response has also been observed in mouse models^15–17^. A mixture of two mouse pancreatic ductal adenocarcinoma (PDAC) cell lines, one giving rise to immune hot and the other to immune cold tumors on its own, resulted in an overall cold tumor, suggesting that coldness may be a dominant phenotype^15^. Another study in a murine melanoma model showed that high-heterogeneity tumors exhibited an inferior CD8 T cell response compared to low-heterogeneity tumors^16^. Building on this, we set out to ask not only how tumor heterogeneity impacts the immune response in the tumor as a whole, but also to interrogate its impact on the spatial organization of the intratumoral immune response.

Our studies employed tumor cell lines derived from carcinogen-induced squamous cell skin carcinomas, induced by dimethylbenzanthracene and 12-O-tetradecanoylphorbol-13-acetate (DMBA/TPA)^18, 19^. The DMBA/TPA skin carcinogenesis model gives rise to tumors that exhibit a physiologically relevant range of mutational burdens^20, 21^ and develop in immune-competent hosts, resulting in tumors with neoantigen and immune profiles that mimic those found in human tumors. We have generated cell lines from DMBA/TPA-induced tumors, which, when implanted into mice, give rise to tumors with reproducible immune phenotypes. Here, we develop a novel model of heterogeneous tumors by implanting a mixture of two such cell lines. Each cell line, labeled with a fluorophore, represents a distinct, trackable population within the tumor. The two cell lines we selected share a common oncogenic driver, *Hras* Q61L, but also each contain over 100 distinct nonsynonymous mutations; their combination thus models a genetically heterogeneous tumor.

Using this novel model system, we here interrogate the mechanisms by which heterogeneity in tumor cells impacts the immune response, the spatial organization of infiltrating immune cells, and the response to ICB therapy. We find, not surprisingly, that the cold tumor population drives an overall cold immune phenotype at a whole tumor level. Further, we observe that our constituent tumor populations form a patchwork pattern within the tumor. Our study reveals that the spatial organization of the tumor cells themselves creates an architectural blueprint that dictates the spatial localization and function of infiltrating immune cells. We observe T cells preferentially accumulating and exhibiting superior effector function in the neighborhood of hot tumor cells, while cold tumor cells recruit an immunosuppressive microenvironment to their immediate vicinity. The presence of the cold tumor cells further mediates resistance to ICB therapy in mixed-population tumors. However, to our surprise, we observe that immunotherapy successfully improves the quality of the immune response in cold tumor regions—but not to a sufficient degree to induce robust tumor regression. Our model system of heterogeneity thus illuminates how heterogeneity in tumor cells impacts both the endogenous anti-tumor immune response and the efficacy of therapy on a highly localized spatial scale.

## Results

### Modeling heterogeneous tumors with trackable tumor populations

We have established a series of squamous cell skin carcinoma cell lines, named the CIT (“carcinogen-induced tumor”) lines, derived from DMBA/TPA-induced skin carcinomas that were initiated in *K5-CreER^T^*^2^-*Confetti* mice on an FVB background^19^. To establish a model system in which to study the impact of tumor heterogeneity on the immune response, we selected and combined two of these cell lines, CIT6 and CIT9, that give rise respectively to tumors exhibiting a high and low frequency of overall T cells, CD4 T cells and CD8 T cells, representing immunologically hot and cold tumors. We introduced fluorescent labels into these tumor cells, taking advantage of CIT lines originating from mice carrying an unactivated *Confetti* cassette. We treated CIT6 and CIT9 with adenoviral Cre recombinase (AdCre), leading to recombination of the *Confetti* allele and stochastic labeling of each cell with one of four fluorescent proteins—yellow, red, cyan and green fluorescent protein (YFP, RFP, CFP and GFP, respectively)^22^. We then sorted YFP^+^ cells from the CIT6 line and RFP^+^ cells from the CIT9 line to establish fluorescently-tagged CIT6-YFP and CIT9-RFP cell lines. For all subsequent experiments, these fluorescently labeled cell lines were implanted into syngeneic *Confetti* mice, carrying an unactivated *Confetti* allele, which recapitulated the expected hot and cold immune phenotypes we observed when unlabeled CIT6 and CIT9 tumor cells were implanted into wild-type FVB mice (**Fig. 1A-D**; Supp Fig. 1).

**Figure 1.**
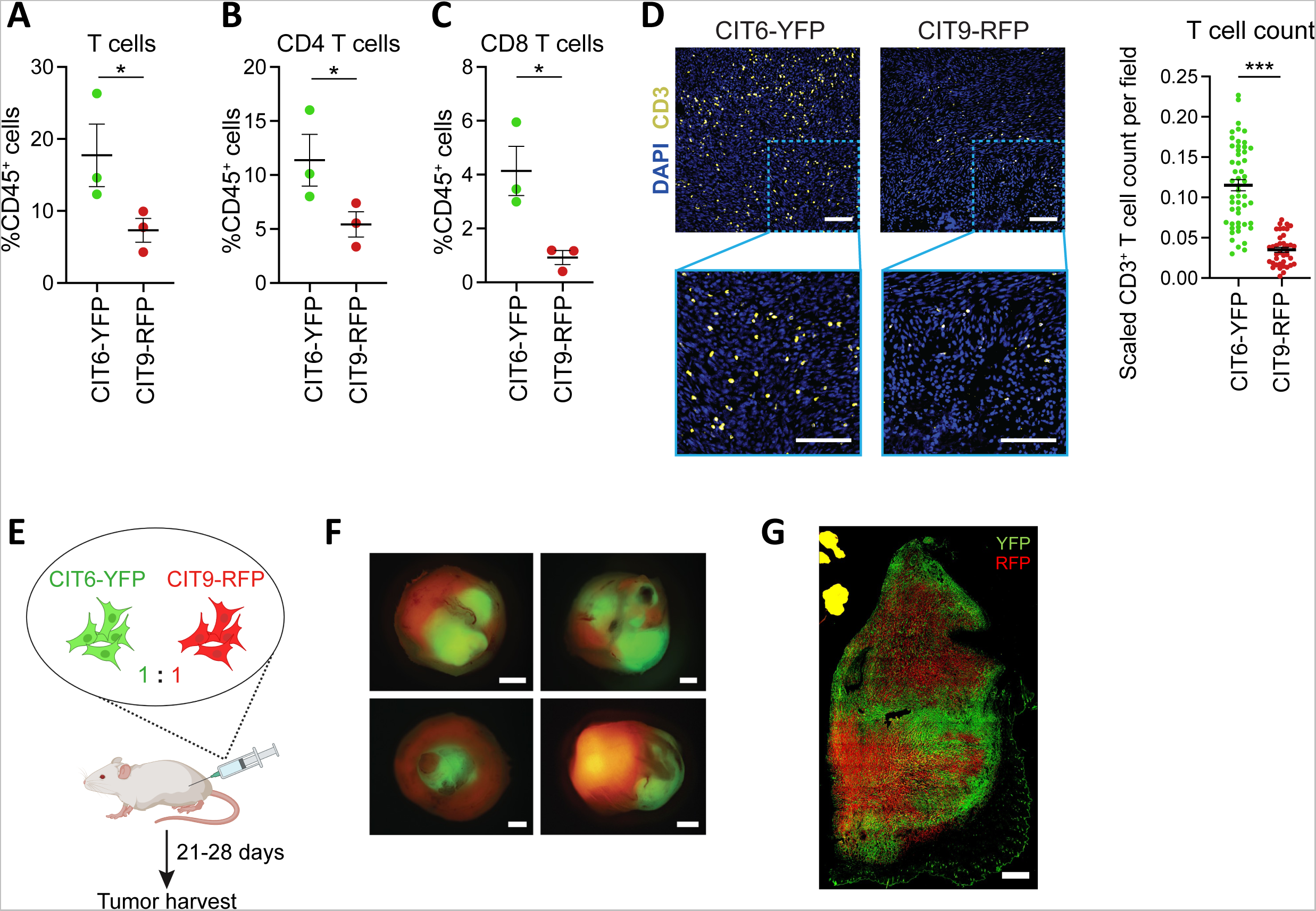
Modeling intratumoral heterogeneity using fluorescently-labeled DMBA/TPA-induced squamous cell skin carcinoma cell lines. **(A-C)** Flow cytometric analysis of T cell infiltration in subcutaneous tumors derived from injection of CIT6-YFP and CIT9-RFP cell lines into mice carrying an unactivated *Confetti* cassette. **(D)** Immunofluorescent staining of CD3^+^ T cells in 8µm cryosections of subcutaneous CIT6-YFP and CIT9-RFP tumors, with DAPI staining cell nuclei, and quantification of T cell count per field Scale bar = 100µm. **(E)** Schematic of mixed-population tumor generation. A 1:1 mixture of CIT6-YFP and CIT9-RFP cell lines subcutaneously was injected into the right hind flank of mice carrying an unactivated *Confetti* cassette, and the resulting tumors were harvested for downstream analyses when they reached 1cm in diameter at 21-28 days post injection. **(F, G)** Whole-tumor images by fluorescent dissecting microscope (F) and a representative image of a tumor cross-section (8µm cryosection) (G) of tumors that were a 1:1 mixture of CIT6-YFP and CIT9-RFP cell lines. Scale bars in (F) and (G) are 2mm and 1mm respectively. All data are a representation of at least 3 independent experiments.

To model heterogeneous tumors, we injected mice subcutaneously with an equal mixture of CIT6-YFP and CIT9-RFP cell lines (6.25×10^4^ cells each, totaling 1.25×10^5^ cells per injection) (**Fig. 1E**). Tumors were assessed for composition by fluorescent stereomicroscope at the time of harvest, when tumors were 1cm in longest diameter (21-28 days). We consistently observed approximately 2/3 of the resulting tumors were comprised of both YFP^+^ and RFP^+^ cells with clearly defined YFP^+^ and RFP^+^ regions, whereas the remaining 1/3 consisted of only RFP^+^ cells in most cases, or only YFP^+^ cells in rare cases (**Fig. 1F**, data not shown). The tumors that contained both YFP^+^ and RFP^+^ cells, termed “mixed-population tumors” herein, were used for downstream analyses. When the equal mixture of CIT6-YFP and CIT9-RFP cells was implanted into immunodeficient NOD/SCID/IL-2Rɣ null (NSG) mice, which lack T, B and NK cells and innate lymphoid cells (ILCs), we observed a higher penetrance of mixed-population tumors, indicating that immune activity against tumor cells was likely responsible for cases where mixed tumors failed to establish (Supp Fig. 2). Inspection of the mixed-population tumors by cryosectioning and fluorescent imaging showed a patchwork of regions that could be classified as being occupied primarily by YFP^+^ cells, primarily by RFP^+^ cells, or by a mixture of both populations (**Fig. 1G**). The fluorescent tags in these cell lines thus successfully enabled spatial tracking and quantification of each tumor population in the mixed-tumor population tumors and enabled us to establish a novel model to study ITH.

### An immunosuppressive phenotype dominates the immune microenvironment of mixed-population tumors

To determine how each constituent tumor population affects the overall immune response in mixed-population tumors, we performed a comprehensive flow cytometric analysis of the immune infiltrates (Supp Fig. 3) found in CIT6-YFP + CIT9-RFP mixed-population tumors and compared this to infiltrates found in each CIT6-YFP and CIT9-RFP single-population tumors. Mixed-population tumors have an overall cold immune phenotype, exhibiting CD4 and CD8 T cell infiltration comparable to CIT9-RFP tumors and significantly lower than that found in CIT6-YFP tumors (**Fig. 2A**). MHC-I proteins were expressed on more than 85% of CIT6-YFP and CIT9-RFP tumor cells and expressed at a similar level (**Fig. 2B**), suggesting that the differences in T cell abundance are not due to the differences in MHC-I expression. MHC-II expression was very low on both CIT6-YFP and CIT9-RFP tumor cells (Supp. Fig. 4A), in line with these tumors being of epithelial origin. Although there was no significant difference in the fraction of CD4 T cells that were regulatory T cells (Tregs; CD4^+^ CD25^+^ Foxp3^+^) or in the CD8 T cell-to-Treg ratio between any of the groups, CIT9-RFP and mixed-population tumors showed a lower CD4 Teff (CD4^+^ CD25^−^ Foxp3^−^)-to-Treg cell ratio than CIT6-YFP tumors, suggesting a skewing in CIT9-RFP and mixed tumors toward an immunosuppressive microenvironment (**Fig. 2C**). Other lymphocytes, specifically B cells and NKp46^+^ lymphocytes (including natural killer (NK) cells and innate lymphoid cells (ILCs)), showed a similar pattern: CIT6-YFP tumors showed significantly higher infiltration by both lymphoid populations than CIT9-RFP tumors, and mixed-population tumors overall resembled CIT9-RFP tumors (although the difference between NKp46^+^ lymphocytes in CIT6-YFP versus mixed tumors was not statistically significant) (**Fig. 2D**). By contrast, CIT9-RFP tumors showed a higher infiltration than CIT6-YFP tumors of F4/80^+^ CD11b^+^ MHC-II^+^ macrophages (**Fig. 2E**) and CD11b^+^ myeloid cells (Supp Fig. 4B), which often harbor immunosuppressive properties in the tumor microenvironment^23, 24^. Mixed-population tumors also had an increase in CD11b^+^ myeloid cells and a trend toward more macrophages compared to CIT6-YFP tumors, again exhibiting more similarity in immune profile to CIT9-RFP tumors (**Fig. 2E**, Supp Fig. 4B). Ly6G^+^ Ly6C^mid^ neutrophils, Ly6G^−^ Ly6C^+^ monocytes, and CD11b^+^ CD11c^+^ MHC-II^+^ F4/80^−^ dendritic cells (DCs), which are conventionally anti-tumoral effector cells, all showed a trend toward lower frequency in both CIT9-RFP and mixed tumors compared to CIT6-YFP tumors, albeit not statistically significant differences (Supp Fig. 4C). Collectively, these data support our designation of CIT6-YFP tumors as pro-inflammatory and immune hot, and of CIT9-RFP tumors as immunosuppressive and immune cold, and further reveal that the immunosuppressive phenotype is dominant in mixed-population tumors.

**Figure 2.**
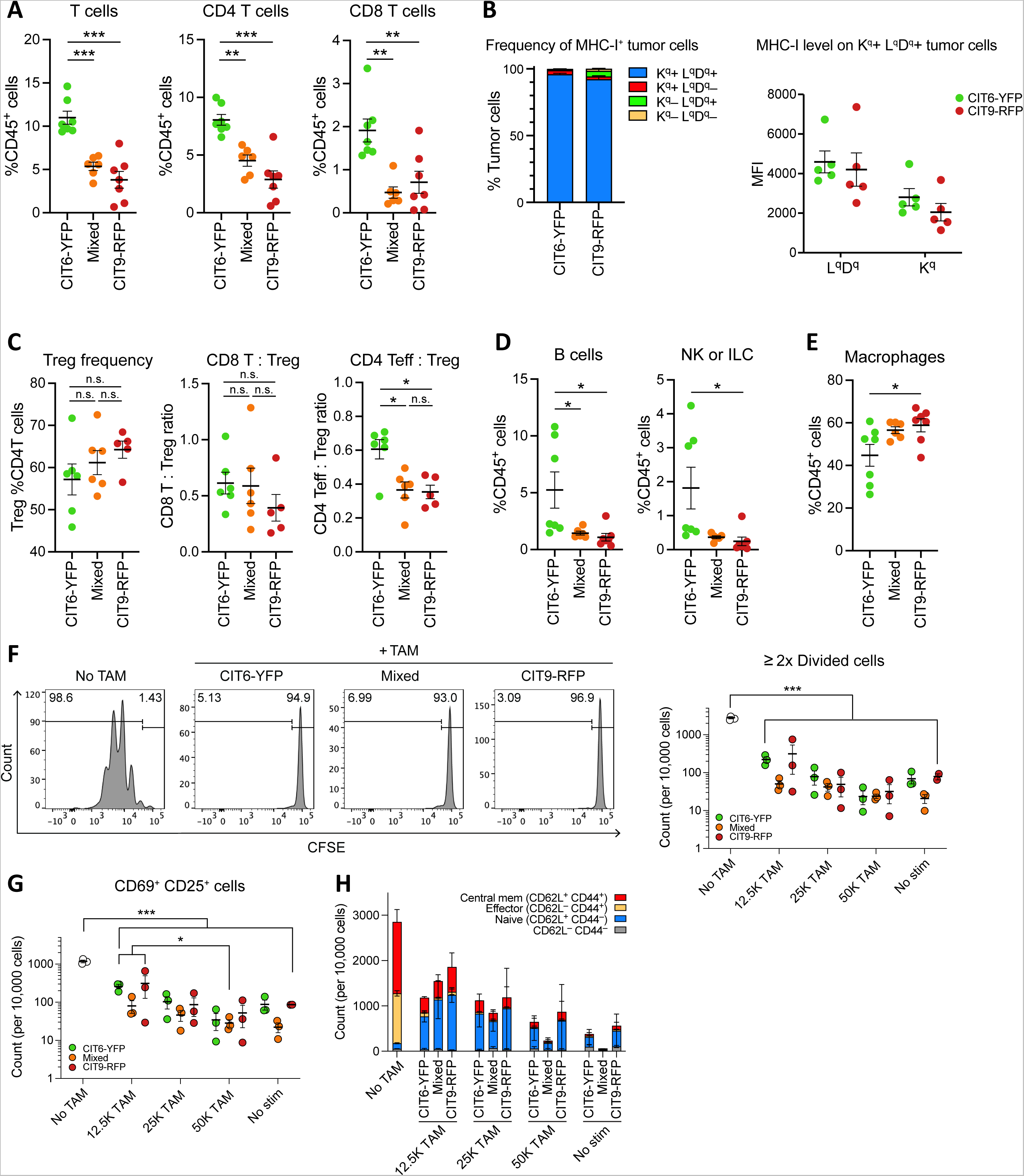
The immunosuppressive tumor population drives an overall immunosuppressive phenotype in mixed-population tumors. **(A-E)** Flow cytometric immune profiling of tumors derived from CIT6-YFP and CIT9-RFP cell lines and mixed-population tumors derived from a 1:1 mixture of the two cell lines, including **(A)** T cells, CD4 T cells and CD8 T cells, **(B)** frequency of tumor cells (CD45^−^ YFP^−^ or CD45^−^ RFP^−^) expressing MHC-I molecules (H2-K^q^ and H2-L^q^D^q^) (left) and median fluorescence intensity (MFI) of the MHC-I molecules on H2-K^q^ and H2-L^q^D^q^ double positive tumor cells (right), **(C)** Treg (Foxp3^+^ CD25^+^) frequency, CD8 T cell-to-Treg ratio, and effector CD4 T cell (Teff, Foxp3^−^ CD25^−^)-to-Treg ratio, **(D)** B cells (B220^+^) and natural killer (NK) or innate lymphoid cell (ILC) (NKp46^+^) lymphocytes and **(E)** macrophages (F4/80^+^ CD11b^+^ MHC-II^+^). Data in (A)-(E) are a representation of at least two independent experiments with ≥5 mice per experiment. **(F-H)** *In vitro* CD8 T cell suppression by macrophages sorted from the tumors. CFSE-stained and CD3/28 beads-stimulated CD8 T cells isolated from naïve *Confetti* mice were co-cultured with tumor-associated macrophages (TAM) at varying ratios (50K T cells and 12.5K, 25K, or 50K TAMs) and analyzed for proliferation via CFSE signal and activation markers. Data are a representation of 2 independent experiments, with 3 biological replicates in each experiment. **(F)** Representative histogram showing the CFSE signal in CD8 T cells, with gates marking divided cells, defined as the cells that have divided more than twice in no TAM condition or 50K T cell + 50K TAM conditions (left) and graph showing the count of divided cells (right). “No TAM”, T cell stimulation control without macrophages; “No stim”, control with no T cell stimulation. **(G)** Count of activated (CD69^+^ CD25^+^) CD8 T cells, **(H)** Count of effector CD8 T cell subsets marked by combinations of CD62L and CD44 expression.

Macrophages are conventionally known to facilitate antigen presentation and assist in T cell activation, but in the context of a tumor, evidence suggests that tumor-associated macrophages (TAMs) are more frequently a mediator of immunosuppression^24^. TAMs are the most abundant immune cells in CIT6-YFP and CIT9-RFP tumors, comprising 30-80% of total CD45^+^ immune cells. Therefore, we sought to determine if F4/80^+^ CD11b^+^ MHC-II^+^ macrophages in our tumors are functionally immunosuppressive. Macrophages were sorted from CIT6-YFP, CIT9-RFP, and mixed-population tumors, and assessed for their capacity to suppress CD8 T cells isolated from naïve, non-tumor bearing syngeneic *Confetti* mice and activated by CD3/CD28 beads. In comparison to CD8 T cells alone, the addition of TAMs significantly suppressed CD8 T cell proliferation, measured as the number of cells that had undergone division (as marked by CFSE dilution) (**Fig. 2F**). The expansion of activated T cells, measured as CD69^+^ CD25^+^ CD8^+^ T cells (early activation), CD69^−^ CD25^+^ CD8^+^ T cells (late activation), or PD-1^+^ CD8^+^ T cells, showed similar results, with the addition of macrophages decreasing the number of activated T cells (**Fig. 2G**, Supp Fig. 4D). Moreover, the expansion of the CD62L^−^ CD44^+^ effector T cell subset and the CD62L^+^ CD44^+^ central memory T cell subset was significantly suppressed in the presence of TAMs (**Fig. 2H**, Supp Fig. 4E). Surprisingly, no difference in immunosuppressive capacity was observed between TAMs derived from CIT6-YFP, CIT9-RFP or mixed-population tumors (**Fig. 2F-H**). Thus, macrophages in CIT6-YFP, CIT9-RFP and mixed-population tumors are equally immunosuppressive, but we conclude that the increased abundance of these macrophages in CIT9-RFP and mixed-population tumors (**Fig. 2E**) may be responsible for creating a more suppressive immune microenvironment in those tumors.

### Spatial organization of tumor populations drives the spatial organization of immune cells

Since the mixed-population tumors were comprised of a patchwork of YFP, mixed, and RFP tumor regions, we next sought to better understand whether the “dominant cold” immune phenotype was uniformly true in all tumor regions. To this end, we first quantified CD3^+^ T cells by immunofluorescent (IF) staining of 8µm sections. We classified each field of view as predominantly occupied by CIT6-YFP cells (“YFP regions”; >60% of tumor cells are YFP^+^), predominantly occupied by CIT9-RFP cells (“RFP regions”; >60% of tumor cells are RFP^+^), or mixed (“mixed regions”; YFP and RFP each represent more than 40% of tumor cells), and then quantified the number of CD3^+^ T cells in the field (**Fig. 3A**). This analysis revealed a significantly higher average count of CD3^+^ T cells in YFP regions compared to RFP regions, with mixed regions showing an intermediate level (**Fig. 3B**). Further, we observed a significant positive correlation between the number of T cells and the fraction of YFP tumor cells (R^2^=0.22, *p*=4.7 × 10^−35^) in each field, and noticed that the fields of view with the highest number of infiltrating T cells were all comprised of >50% YFP^+^ cells (**Fig. 3C**). These data suggest that T cells preferentially localize to the regions occupied by CIT6-YFP tumor cells in mixed-population tumors.

**Figure 3.**
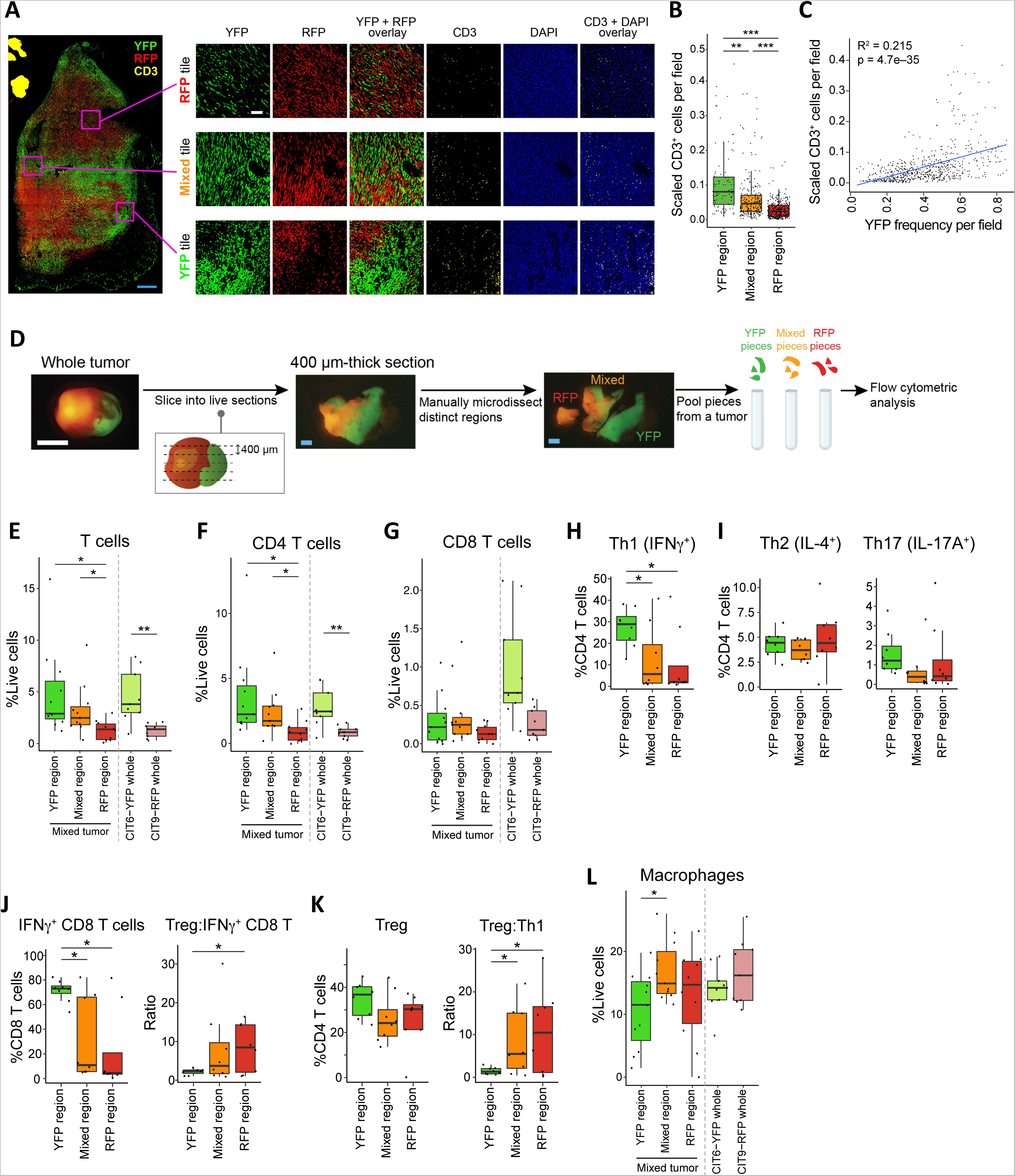
Spatial organization of tumor populations drives the spatial organization of immune cells. **(A)** Representative cross-section images showing examples of a YFP, mixed and RFP region of a mixed-population tumor and CD3 stain. **(B)** Quantification of CD3^+^ T cells per field, plotted for each region. **(C)** Correlation of YFP^+^ cell fraction and CD3^+^ T cell frequency in each field. CD3^+^ T cell count was scaled to DAPI^+^ cell count in each field to normalize the difference in cell density between fields. **(D)** Schematic of immune cell profiling in YFP, mixed and RFP regions of mixed-population tumors. Each mixed-population tumor was sliced into five to ten 400µm-thick live sections, and each section was manually microdissected into YFP, mixed, and RFP regions using a surgical scalpel under a fluorescent dissecting microscope. All YFP pieces, all mixed pieces, and all RFP pieces from the same tumor were pooled for immune profiling by flow cytometry. White scale bar = 5mm, blue scale bar = 1mm. **(E-L)** Frequency of T cell subsets, ratio of Treg to IFN*γ*^+^ CD8 T cells or Th1 cells, and frequency of macrophages in YFP, mixed and RFP regions. Grey dotted lines in panels (E), (F), (G) and (L) separate data of mixed-population tumors from the data of single-population tumors. All data are a representation of two independent experiments, with ≥3 mice per experiment (B,C) or ≥6 mice per experiment (E-L).

We next turned to flow cytometry-based analysis of the immune cells in each YFP, mixed, and RFP regions to gain a higher dimensional profile of the immune response in each region. Mixed population tumors were embedded into agarose gel and each tumor was sliced into five to ten 400µm-thick live sections. Each section was manually microdissected into YFP, mixed, and RFP regions using a surgical scalpel under a fluorescent dissecting microscope. All YFP pieces, all mixed pieces, and all RFP pieces from the same tumor were pooled and digested into a single cell suspension for immune profiling by flow cytometry (**Fig. 3D**). Mirroring our findings by IF, we found a higher frequency of total CD3^+^ T cells in YFP regions than in RFP regions, with mixed regions showing intermediate infiltration (**Fig. 3E**). This localization pattern was also seen in, and was driven by, CD4 T cells. The abundance of total T cells and CD4 T cells in YFP regions of mixed-population tumors was comparable to that in single-population CIT6-YFP tumors, and similarly, the abundance of total and CD4 T cells in RFP regions of mixed tumors was comparable to that in single-population CIT9-RFP tumors (**Fig. 3F**). By contrast, CD8 T cells were at low abundance in all tumor regions, including YFP regions, and were found at levels resembling those in single-population CIT9-RFP tumors (**Fig. 3G**). A more detailed analysis of T cell function showed that nearly one-third of the CD4 T cells in YFP regions were Th1 cells (CD4^+^ IFN*γ*^+^), whereas only ∼2% of CD4 T cells in RFP regions were Th1 cells (median frequency of Th1 cells in YFP regions = 28.9%, RFP regions = 1.99%) (**Fig. 3H**). Th2 (CD4^+^ IL-4^+^) and Th17 (CD4^+^ IL-17^+^) cells were equally rare in all regions (**Fig. 3I**). Even more strikingly, despite their low abundance, CD8 T cells showed increased effector function in YFP regions: over 70% of CD8 T cells in YFP regions showed capacity to produce IFN*γ*, whereas less than 10% of CD8 T cells in mixed and RFP regions showed the same (**Fig. 3J**). There was no apparent difference in Treg (Foxp3^+^ CD25^+^) cell frequency between regions, but the ratio of Treg:Th1 cells and Treg: IFN*γ*^+^ CD8 T cells was higher in RFP regions compared to YFP and mixed regions, suggestive of an overall more immunosuppressive microenvironment in RFP regions (**Fig. 3J, K**). In addition, macrophages (F4/80^+^ CD11b^+^ MHC-II^+^) were significantly enriched in mixed regions compared to YFP regions, and also trended toward higher enrichment in RFP regions (**Fig. 3L**).

Collectively, these data show that local tumor cells themselves play a significant role in driving the immune microenvironment in their vicinity. Moreover, the observation that mixed regions look similar to RFP regions—in terms of effector T cell infiltration and macrophage infiltration—suggests that in regions in which both CIT6-YFP and CIT9-RFP tumor cells co-exist, cold CIT9-RFP tumor cells exert dominance over the local immune infiltrate. The “dominant coldness” of CIT9-RFP is also seen in the global suppression of CD8 T cell infiltration in all regions. However, CD8 T cells that managed to infiltrate YFP regions show a highly active phenotype.

### CD206^Hi^ macrophages and lack of neutrophils and monocytes mediate an immunosuppressive TME in RFP regions

To characterize immune cell spatial patterning more deeply, we employed ZipSeq spatial transcriptomics^25^ to CIT6-YFP + CIT9-RFP mixed-population tumors. ZipSeq allows immune cells within user-defined regions of interest in live tissue sections to be tagged with distinct nucleotide sequences (akin to assigning each region of interest a unique zip code), making use of photocaged oligonucleotides and precise light exposure. Using this method, immune cells from YFP, RFP and mixed regions of mixed population tumors (n=2) were tagged with distinct “zip code” barcodes, and sections were subsequently disaggregated and subjected to single cell RNA sequencing (scRNA-seq) (**Fig. 4A**). Immune cells were subjected to UMAP clustering, and cluster identities were assigned based on expression of well-established markers (**Fig. 4B**, Supp Fig. 5A, B). Using ZipSeq barcodes, we mapped each immune cell back to the region it originated from. In line with our flow cytometry analyses, we observed a higher infiltration of T cells in YFP regions compared to RFP regions (**Fig. 4C, D**), and also saw that myeloid cells comprised a majority of the immune cells (**Fig. 4B**). In particular, we observed three large macrophage clusters, CD206^Hi^ macrophages, MHC-II^Hi^ macrophages, and Arg1^Hi^ macrophages (**Fig. 4B**, Supp Table 1). These three clusters were all enriched in RFP regions (**Fig. 4E**). By contrast, neutrophils and monocytes were observed at lower abundance in RFP regions than YFP regions (**Fig. 4F, G**). We carried out ZipSeq single cell spatial transcriptomics on a second mixed-population tumor, and found similar results (Supp Fig. 6). Myeloid cells comprised a majority of the immune cells (Supp Fig. 6A-C) and T cells were enriched in YFP regions (Supp Fig. 6D, E). Notably, out of seven macrophage clusters in the second tumor, six of them were enriched in RFP regions compared to YFP regions (Supp Fig. 6F). Moreover, the second tumor showed enrichment of neutrophils and monocytes in YFP regions (Supp Fig. 6G, H). Pathway analysis with EnrichR, using genes enriched in each cluster and the GO Biological Process 2021 repository, suggested neutrophils had pro-inflammatory properties, as seen by neutrophil activation-related genes (**Fig. 4F**, Supp Fig. 6G). Monocytes also exhibited evidence of activation, with a particularly clear pro-inflammatory signature in the second tumor (**Fig. 4G**, Supp Fig. 6H). Together, the combination of an enrichment of immunosuppressive macrophages and a de-enrichment of inflammatory monocytes and neutrophils point to a myeloid cell-driven immunosuppressive TME in RFP regions.

**Figure 4.**
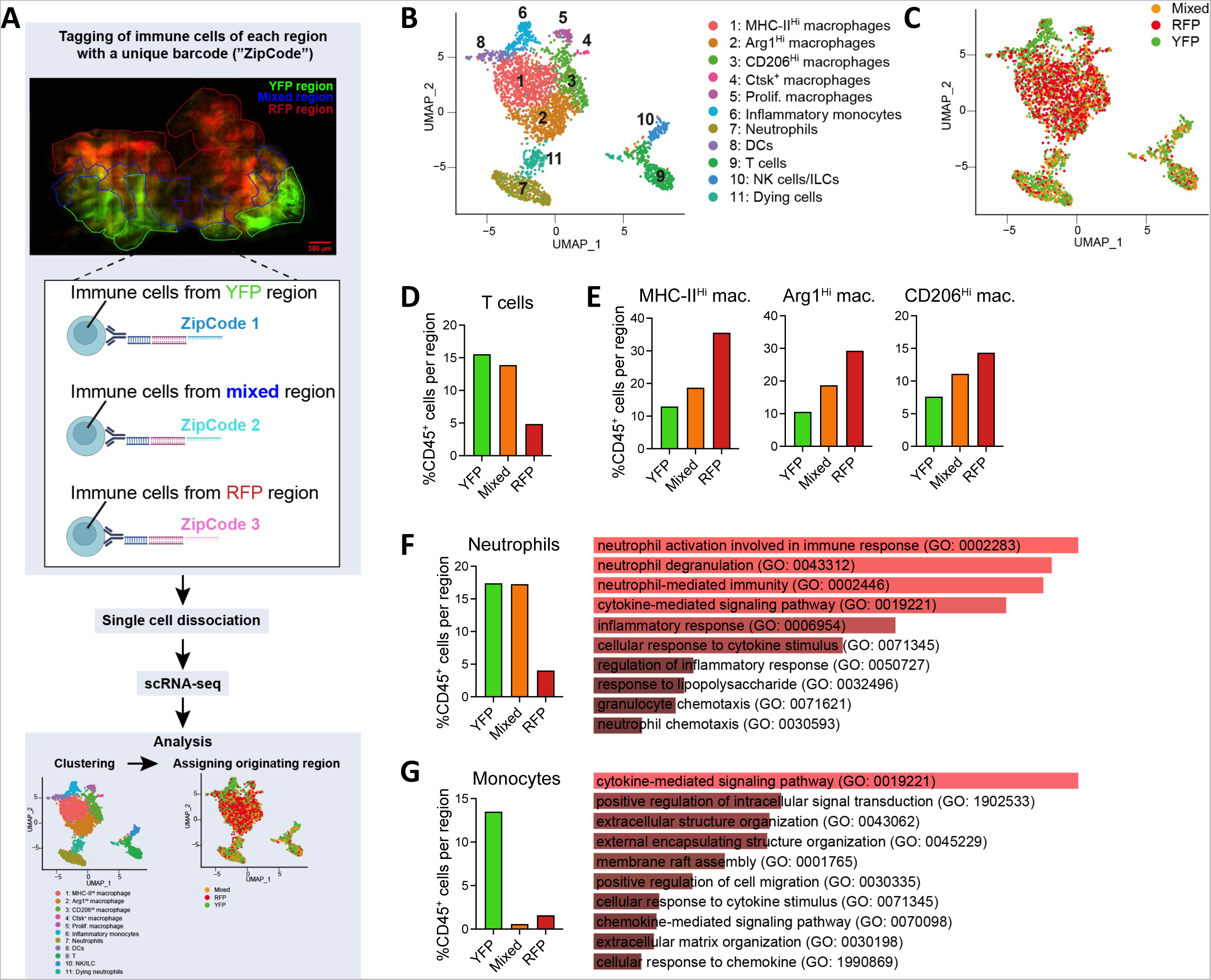
CD206^Hi^ macrophages and lack of neutrophils and monocytes mediate an immunosuppressive TME in RFP regions. **(A)** Schematic of ZipSeq spatial transcriptomic analysis of mixed-population tumors. Composite stitched image of a 200µm-thick live section marked with YFP regions (encircled with green lines), RFP regions (red lines) and mixed regions (blue lines) is shown. YFP, RFP and mixed regions were illuminated individually to allow for the binding of a unique barcode (“zipcode”) to a photocaged oligo-CD45 antibody complex. ZipCode-tagged immune cells were analyzed by scRNAseq. **(B)** Uniform manifold approximation and projection (UMAP) representation of zipcode-labeled cells with cluster overlay. *n* = 3,238 cells, *n_YFP_* = 1,236, *n_RFP_* = 1,318, *n_Mixed_* = 684. **(C)** UMAP representation of zipcode-labeled cells with zipcode identity overlaid. **(D)** Abundance of cells belonging to the T cell cluster (cluster 9), calculated as percentage of total immune cells in each region. **(E)** Abundance of cells belonging to MHC-II^Hi^, Arg1^Hi^ and CD206^Hi^ macrophage clusters (clusters 1, 2, and 3 respectively), calculated as percentage of total immune cells in each region. **(F)** Percentage of cells belonging to neutrophil cluster (cluster 7) in each region and pathway analysis showing the gene families enriched in the neutrophil cluster based on the top 250 differentially expressed genes. **(G)** Percentage of cells belonging to monocyte cluster (cluster 6) and pathway analysis of genes enriched in the monocyte cluster based on the top 31 differentially expressed genes.

### Chemokine CX3CL1 is a mediator of the CIT9-RFP-driven immunosuppressive TME

To identify the drivers behind the striking spatial organization of myeloid cell infiltrates in our mixed tumors, we next assessed the protein expression of a total of 44 cytokines and chemokines using the lysates from CIT6-YFP and CIT9-RFP tumors. Cytokines and chemokines are major mediators of immune cell localization and function, thus we hypothesized differences in their expression between CIT6-YFP and CIT9-RFP might explain the differences we saw in myeloid infiltrate. We found the chemokine CX3CL1 was significantly enriched in CIT9-RFP tumors compared to CIT6-YFP tumors (**Fig. 5A**, Supp Fig. 7A). CX3CL1 is a chemokine with reported pro-tumor properties, capable of directly promoting tumor cell proliferation and migration as well as of attracting various immune cell types^26, 27^. To test the hypothesis that CX3CL1 plays a role in driving the immunosuppressive microenvironment in our squamous cell carcinomas, we generated a CIT6-YFP cell line overexpressing *Cx3cl1* and analyzed the immune profile of these tumors. The abundance of macrophages was increased in Cx3cl1-overexpressing tumors compared to control tumors (**Fig. 5B**). We next asked if it was a particular macrophage subset(s) that showed increased abundance, and for this analysis divided macrophages into the 3 main clusters we identified in our ZipSeq analysis: CD206^Hi^ macrophages (CD206^Hi^ MHC-II^Low^), MHC-II^Hi^ macrophages (CD206^Mid^ MHC-II^Hi^ Arg1^−^) and Arg1^Hi^ macrophages (CD206^Mid^ MHC-II^Hi^ Arg1^+^) (Supp Fig. 7B). We found that CD206^Hi^ macrophages were increased in frequency in CX3CL1-overexpressing CIT6-YFP tumors compared to control tumors, while the other two macrophage subsets showed no difference in frequency (**Fig. 5C**). In line with this, expression of CX3CR1, a cognate receptor of CX3CL1, was the highest on macrophages among all immune cell types (Supp Fig. 7C), and highest on CD206^Hi^ macrophages among the three macrophage subsets (**Fig. 5D**). Additionally, we observed a lower abundance of neutrophils (Ly6G^+^) and monocytes (Ly6C^+^) in CX3CL1-overexpressing CIT6-YFP tumors (**Fig. 5E**). Other immune cell types including B cells, CD8 T cells and DCs showed no difference in frequency (Supp Fig. 7D).

**Figure 5.**
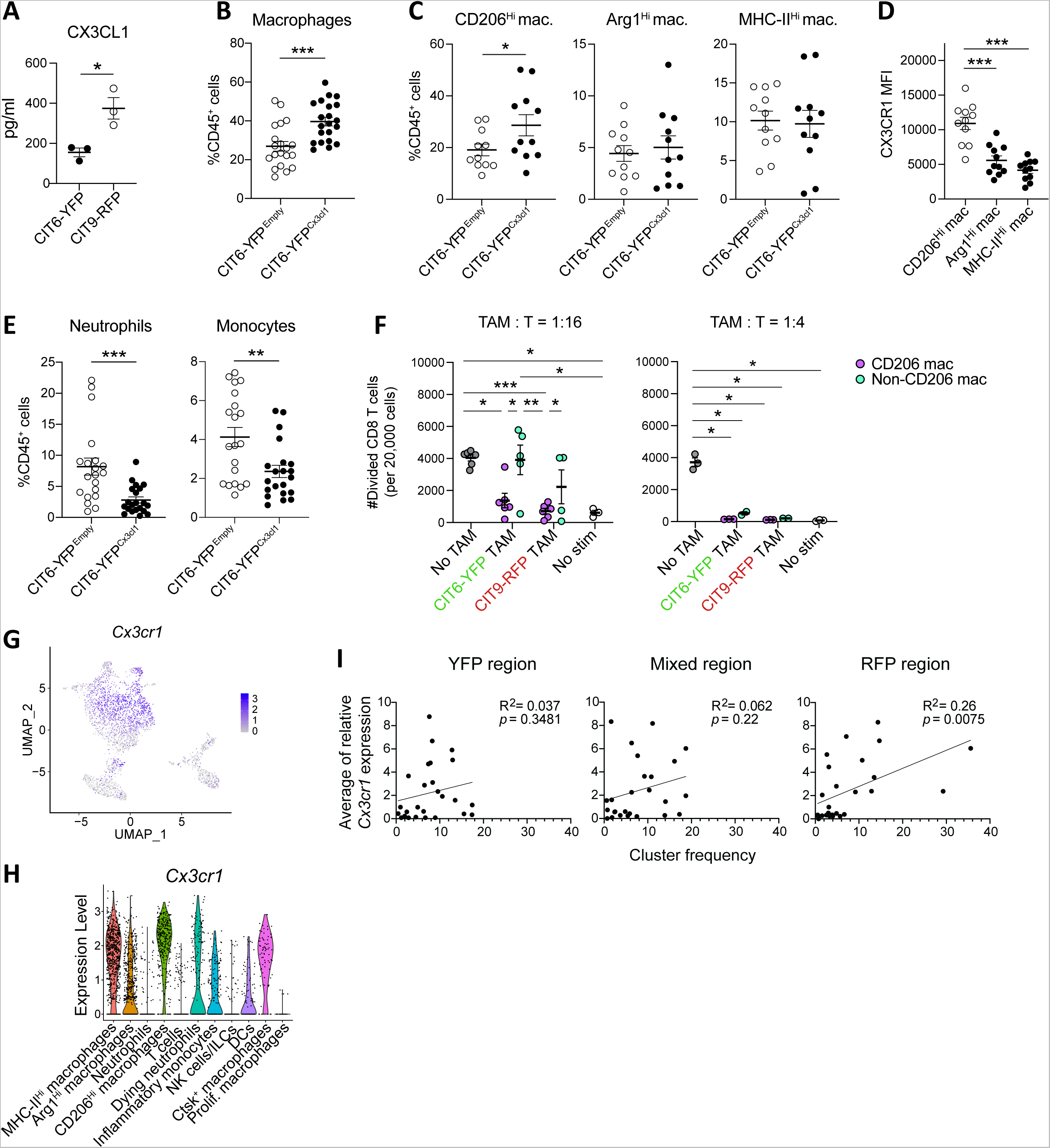
Chemokine Cx3cl1 mediates the CIT9-RFP-driven immunosuppressive TME. **(A)** ELISA-based quantification of CX3CL1 using tumor lysates of CIT6-YFP and CIT9-RFP single-population tumors (n=3 per cell line) **(B, C)** Infiltration of total macrophage (B) and macrophage subsets (C) in CIT6-YFP tumors overexpressing CX3CL1 (CIT6-YFP^Cx3cl1^) and empty vector (CIT6-YFP^Empty^). **(D)** Expression of CX3CR1 on macrophage subsets in CIT6-YFP^Cx3cl1^ tumors, measured as MFI from flow cytometric analysis. **(E)** Frequency of neutrophils and monocytes in CX3CL1-overexpressing CIT6-YFP tumors. Data in (B)-(E) are representation or combination of two independent experiments with *≥*10 mice per experiment. **(F)** CD8 T cell suppression assay using CD206^Hi^ macrophages and a pool of Arg1^Hi^ macrophages and MHC-II^Hi^ macrophages (Non-CD206^Hi^ TAMs) isolated from CIT6-YFP and CIT9-RFP single-population tumors. Representation of two independent experiments with at least 2 mice per arm. **(G)** Feature plot overlaid on UMAP representation for *Cx3cr1* gene from the tumor analyzed by ZipSeq in Fig. 4. **(H)** Violin plot showing *Cx3cr1* distribution based on cluster identity. **(I)** Correlation between an average relative *Cx3cr1* transcript level of each cluster and the percentage of cells belonging to the cluster. Combined analysis of two tumors analyzed in Fig. 4 and Supp Fig. 6.

We next asked whether CD206^Hi^ macrophages, as the main macrophage subtype increased by CX3CL1 overexpression, had a greater capacity to suppress T cells than non-CD206^Hi^ macrophages (which include both MHC-II^Hi^ and Arg1^Hi^ macrophages). We isolated CD206^Hi^ and non-CD206^Hi^ macrophages from CIT9-RFP and CIT6-YFP tumors by flow cytometry-based sorting (Supp Fig. 7B) and tested for their ability to suppress CD8 T cell proliferation using an *in vitro* co-culture assay. We found that CD206^Hi^ macrophages could suppress T cell proliferation and activation even at a low macrophage-to-CD8 T cell ratio (1:16), whereas non-CD206^Hi^ macrophages could suppress CD8 T cells only at a higher co-culture ratio (1:4) (**Fig. 5F**, Supp Fig. 7E). Thus, while both CD206^Hi^ and non-CD206^Hi^ macrophages exhibited suppressive capacity to some extent, the CD206^Hi^ macrophage subset in our tumors has a particularly strong ability to suppress T cell proliferation.

Because we also observed an enrichment of CD206^Hi^ macrophages and lower abundance of neutrophils and monocytes in RFP regions of mixed-population tumors in our ZipSeq data (**Fig. 4**), we hypothesized that the CX3CL1-CX3CR1 axis plays a role in driving the immunosuppressive microenvironment in these regions. There were few reads of *Cx3cl1* in the ZipSeq data (data not shown) likely due to limitations associated with single cell RNAseq. Therefore, to test this hypothesis, we assessed the correlation between the *Cx3cr1* expression level in immune cells and their abundance in RFP regions. *Cx3cr1* transcript level was high in clusters that were enriched in RFP regions (**Fig. 5G** and **Fig. 4C**; Supp Fig. 8A and Supp Fig. 6D), and was the highest on the CD206^Hi^ macrophage cluster (**Fig. 5H**, Supp Fig. 8B). Moreover, there was a positive correlation across clusters between the abundance of a cluster and its average *Cx3cr1* transcript expression level in RFP regions, but not in YFP or mixed regions (**Fig. 5I**). Collectively, these data indicate that CX3CL1 plays a role in driving local myeloid cell abundance and enriching immunosuppressive CD206^Hi^ macrophages in RFP regions of mixed-population tumors.

### CX3CL1 is a mediator of a cold TME in human cancers

To assess the role of CX3CL1 in shaping the TME and myeloid cell abundance in human cancers, we turned to a previously published pan-cancer human dataset containing transcriptomic profiles of whole tumors and of individual immune compartments (e.g., myeloid compartment) from 364 individual tumors across 12 cancer types^28^. Previous analysis identified 12 recurrent immune infiltration patterns across these 364 tumors, denoted “immune archetypes”, spanning a range from immune rich to immune poor TMEs. Looking at T cell abundance, 7 of these archetypes could be classified as immune hot and 5 of them as immune cold, corresponding with high and low T cell infiltrates based on T cell transcriptomic scores (**Fig. 6A**). We assessed whole-tumor *CX3CL1* RNA expression in each archetype, and found that, strikingly, *CX3CL1* levels were highest in 3 of 5 cold archetypes (archetypes 9, 10 and 12) and comparatively low in all hot archetypes (**Fig. 6B**). Next, we examined the macrophage:monocyte ratio across all human archetypes, using transcriptomic scores for macrophages and monocytes based on RNAseq data from the myeloid compartment of each tumor.^28^ We observed a bias in the myeloid compartment toward macrophages (log2 of the macrophage:monocyte ratio > 0) in the same 3 cold human archetypes that showed high *CX3CL1* (archetypes 9, 10, and 12, **Fig. 6C**). Using a gene signature based on the CD206^hi^ cluster in our ZipSeq dataset, we also asked if CD206^hi^ macrophages were enriched in *CX3CL1*-high human immune archetypes. We found that, among cold archetypes, the macrophage compartment showed a high CD206^hi^ macrophage score in 2 archetypes, both of which were *CX3CL1*-high (archetypes 10 and 12, **Fig. 6D**). Finally, we turned back to our CX3CL1-overexpressing mouse tumors. We saw that CX3CL1 overexpression led to an increase in both the macrophage:monocyte ratio and the CD206^hi^ macrophage: non-CD206^hi^ macrophage ratio in CX3CL1-overexpressing CIT6-YFP tumors compared to controls (**Fig. 6E**). Of note, we also analyzed the 7 hot human archetypes that show low *CX3CL1* expression, and found a range of macrophage:monocyte ratios and CD206^hi^ macrophage scores (Supp. Fig. 9), suggesting that additional and likely different factors shape these ratios in hot, CX3CL1-low tumors. Collectively, these data show that CX3CL1 is associated with cold TMEs enriched in CD206^hi^ macrophages not only in our mouse model, but also across multiple human cancers.

**Figure 6.**
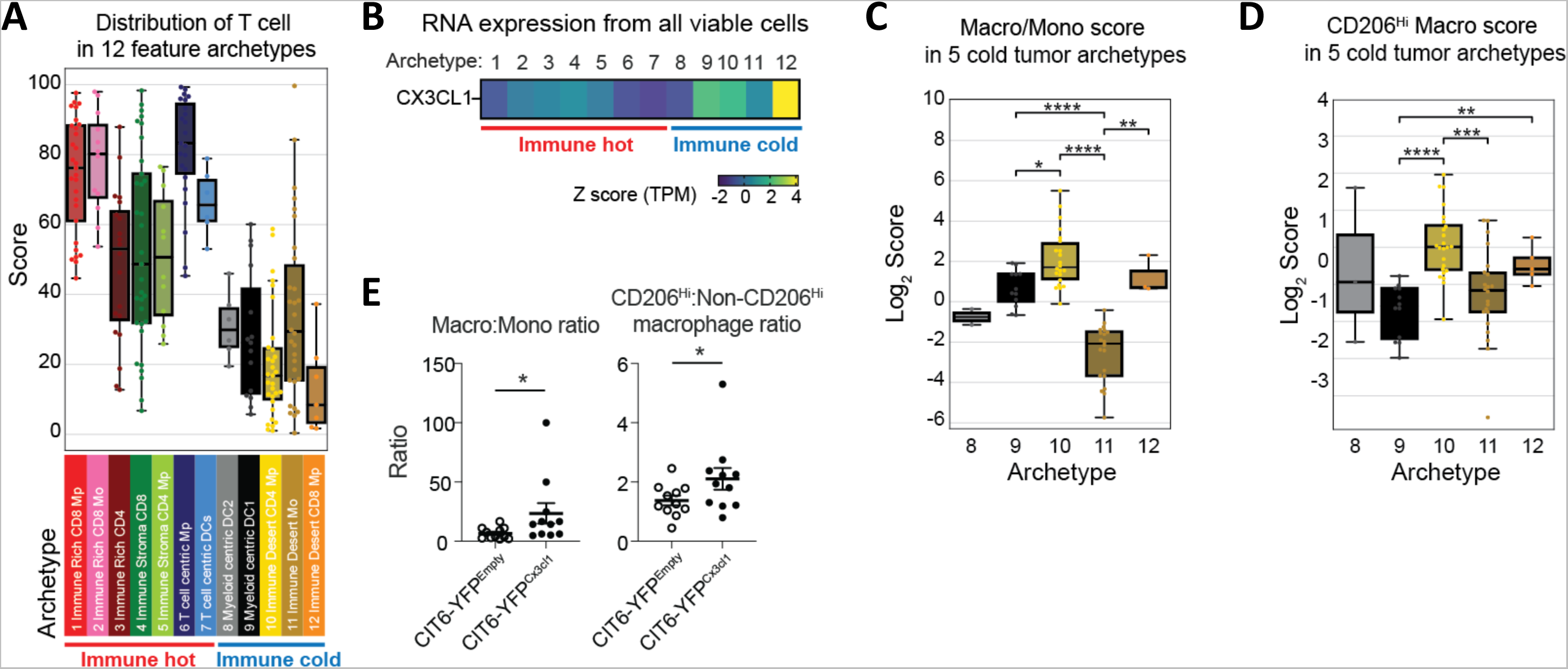
CX3CL1 is a mediator of a cold TME in human cancers. **(A-D)** Analysis of immune cells and *CX3CL1* expression using a previously published dataset of 364 human tumors from 12 cancer types. Computational clustering of tumors using flow cytometry data and transcriptomic data enabled identification of 12 dominant patterns of immune composition across various cancers, referred to as immune archetypes (ref. 28). RNAseq data was generated separately for all viable cells and for sorted mononuclear phagocytes. Prior to RNA sequencing, cells were sorted by flow cytometry from untreated, freshly resected human tumors. **(A)** T cell score of each tumor, based on RNAseq of all viable cells, grouped on the x-axis by immune archetype. **(B)** Heatmap of average *CX3CL1* RNA expression, based on RNAseq of all viable cells, for each immune archetype. **(C)** Ratio of macrophage score and monocyte score, based on RNAseq of mononuclear phagocytes, for tumors in the 5 cold immune archetypes. **(D)** CD206^Hi^ macrophage score, based on RNAseq of mononuclear phagocytes, for tumors in the 5 immune cold archetypes. **(E)** Ratio of macrophages to monocytes (left) and CD206^Hi^ macrophages to non-CD206^Hi^ macrophages (right) detected in mouse CIT6-YFP tumors overexpressing *Cx3cl1* or controls.

### An immunosuppressive tumor population drives resistance of mixed-population tumors to anti-PD-1 blocking and CD40 agonistic antibody combination treatment

Having shown that CIT9-RFP tumor cells in mixed-population heterogeneous tumors exert a dominant immunosuppressive effect and drive spatial organization within the tumor, we next sought to determine if the presence of CIT9-RFP tumor cells impacts the response of mixed-population tumors to immune checkpoint blockade (ICB) therapy. In particular, as an immunosuppressive tumor microenvironment is linked to resistance to ICB therapy, we asked if CIT9-RFP tumor cells make otherwise therapy-sensitive CIT6-YFP tumor cells resistant. We tested multiple ICB combinations on our tumors, and we selected a regimen combining PD-1 blockade and CD40 agonist antibodies, a combination that is currently in clinical trials for multiple tumor types. The combination treatment was administered in two doses at a 3-day interval when the CIT6 and CIT9 tumors reached 5mm in diameter (**Fig. 7A**) and induced 50% of CIT6-YFP tumors (5 out of 10 tumors) to regress completely, with 2 tumors eventually growing back after 5-17 days (**Fig. 7B**). In contrast, only 9% (1 out of 11) of CIT9-RFP tumors showed a decrease in tumor volume after therapy, and none achieved a complete regression (**Fig. 7B**). Furthermore, the increase in the average volume of CIT6-YFP tumors over time was blunted by the combination treatment in comparison to isotype control treatment, whereas CIT9-RFP tumors showed similar growth kinetics in both combination treatment and isotype control arms (**Fig. 7B**). This coincided with prolonged overall survival of the mice harboring CIT6-YFP tumors, but not CIT9-RFP tumors, after treatment with the combination therapy (**Fig. 7C**). Thus, the combination of anti-PD-1 blocking and CD40 agonistic antibodies mimics prototypical patient outcomes to immunotherapy where immune hot tumors respond and immune cold tumors do not. Further, this combination showed the clearest dichotomy in responses between hot and cold tumors among the regimens tested, and so we chose to proceed with using it to investigate how mixed-population tumors would respond.

**Figure 7.**
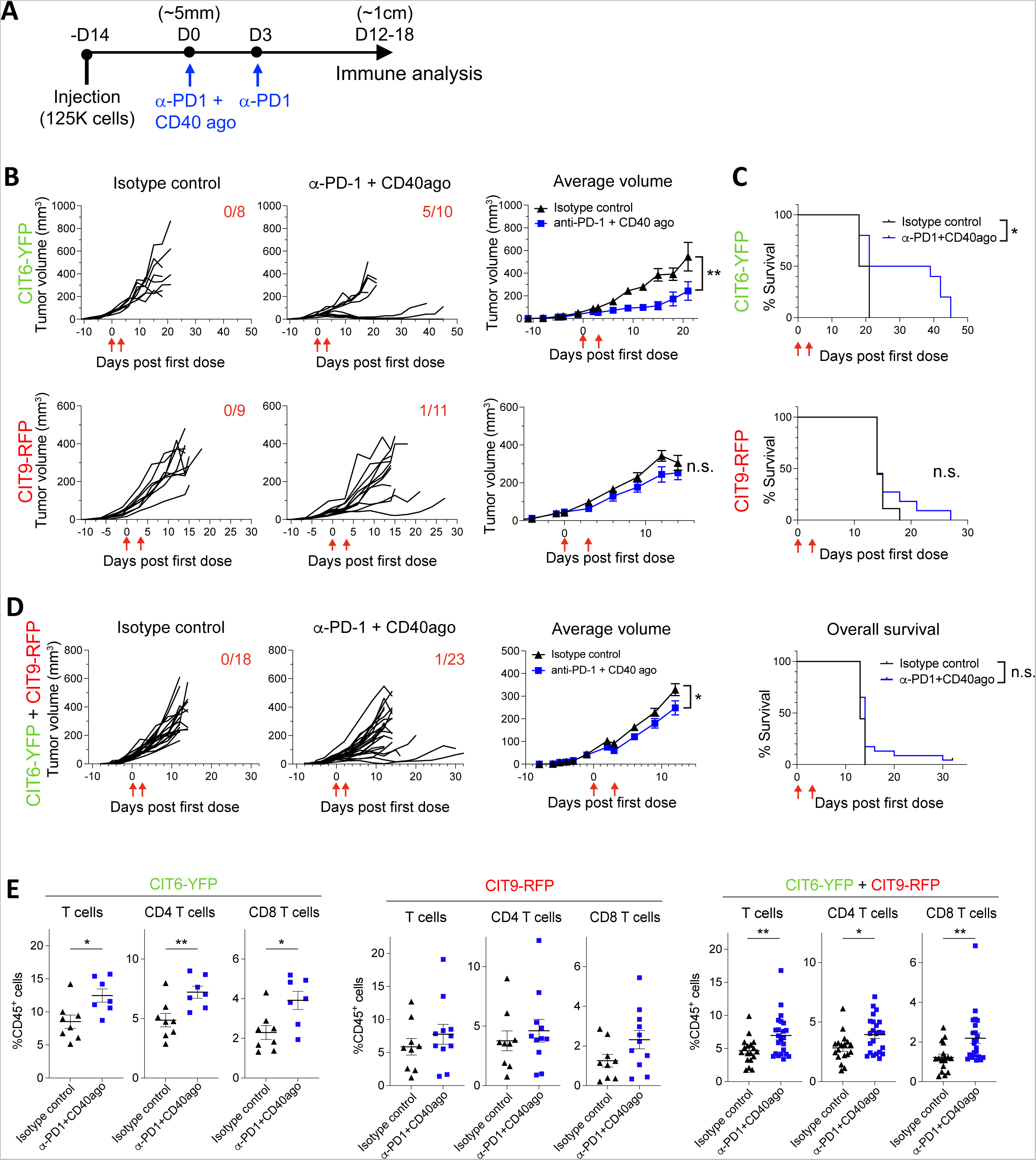
An immunosuppressive tumor population drives resistance of mixed-population tumors to anti-PD-1 blocking and CD40 agonistic antibody combination treatment. **(A)** Dosing schedule of anti-PD-1 and CD40 agonist treatment. **(B)** Tumor growth curves of CIT6-YFP and CIT9-RFP tumors showing tumor volume of individual tumors and average volume of isotype control and anti-PD-1+CD40 agonist combination treatment arms. Red arrows under the x axis shows the timing of treatments (day 0 and 3), fraction shown on the top right corner of individual tumor volume plots shows the number of mice whose tumors completely regressed post-treatment. **(C)** Kaplan-Meier curves comparing the survival of mice from each treatment arm. Statistical test was performed using log-rank test. **(D)** Tumor growth and survival of mice harboring tumors derived from a 1:1 mixture of CIT6-YFP and CIT9-RFP cell lines. **(E)** Frequency of tumor-infiltrating T cells post-treatment in CIT6-YFP, CIT9-RFP, and 1:1 mixture-derived tumors. All experiments were performed with at least 8 mice per treatment arm.

We then assessed the response of tumors derived from a 1:1 mixture of CIT6-YFP and CIT9-RFP cell lines to the anti-PD-1 blocking and CD40 agonist combination treatment. The combination therapy achieved complete regression (CR) in 1 out of 23 mice (4.3%) but the tumor quickly grew back (**Fig. 7D**). The CR rate of mixture-derived tumors was higher than CIT9-RFP single-population tumors (0% CR), but lower than CIT6-YFP tumors (50% CR) (**Fig. 7B, D**). The average tumor volume was slightly smaller in the combination treatment group (**Fig. 7D**), although this did not correlate to an extension of overall survival (**Fig. 7D**), suggesting therapy had only a modest effect in these mixed tumors.

Immune profile analysis of tumors at the time of harvest (10mm in diameter) revealed that CD4 and CD8 T cell infiltration was increased in both CIT6-YFP tumors and mixed tumors treated with the anti-PD-1 + anti-CD40 agonist combination, and that this influx was of similar magnitude in both (**Fig. 7E**). By contrast, CIT9-RFP tumors did not show a change in T cell infiltration after treatment (**Fig. 7E**). However, despite the influx of T cells into mixed tumors, this influx did not correspond to the same level of treatment efficacy seen in CIT6-YFP tumors. Collectively, these data suggest that the presence of CIT9-RFP tumor cells dampens the response of mixed tumors to the combination treatment, and also reveals an improved influx of T cells into the tumor after treatment is not necessarily a marker of a successful response in the context of a heterogeneous tumor.

### Spatial immune infiltration patterns persist in mixed-population tumors following anti-PD-1 blockade and CD40 agonist combination treatment

Given our previous data showing that RFP regions (and to a lesser extent, mixed regions) of mixed-population tumors were spatially organized pockets of immune coldness, we sought to test the hypothesis that the immunotherapy combination treatment was acting successfully in YFP regions, but failing to improve the immune response in RFP regions, leading to overall failure. We applied our live tumor slice microdissection technique (**Fig. 3D**) to mixed-population tumors treated with either anti-PD-1 blockade + CD40 agonist or control antibodies. The immune infiltration profiles of YFP, RFP and mixed regions were analyzed 6 days after the initial treatment dose, at which point tumors have not started to shrink yet (**Fig. 7B and D**). We focused our analysis on T cells, as we had observed that T cell infiltration was increased by the combination treatment in mixed population tumors (**Fig. 7E**).

At the 6-day timepoint, the frequencies of overall CD4 and CD8 T cells did not differ between the isotype control arm and the combination treatment arm in any of the three regions (Supp Fig. 10). However, the presence of Th1 cells was dramatically increased in RFP and mixed regions by the combination treatment compared to isotype control treatment (**Fig. 8A**). There was also a trend toward increased Th1 cells in YFP regions, but it did not reach statistical significance. The treatment-induced change in abundance was unique to Th1 cells, as other CD4 T cell subsets including Treg, Th2 and Th17 cells showed a similar frequency between isotype control and combination treatment arms (**Fig. 8B, C**). The changes in Th1 cell abundance resulted in a favorable increase in the Th1:Treg ratio in mixed and RFP regions of the combination therapy-treated tumors (**Fig. 8D**). The functional impact on the CD8 T cell compartment was directionally the same, but much less dramatic. The frequency of IFN*γ*^+^ CD8 T cells was modestly increased by the combination therapy in RFP regions, but not in mixed regions (**Fig. 8E**), and the IFN*γ*^+^ CD8 T cell to Treg ratio showed a trend of improvement in mixed and RFP regions in response to the treatment (**Fig. 8F**). However, because the abundance of CD8 T cells remained low in all regions (as previously observed in mixed-populations tumors, Fig. 3G), these improvements in CD8 T function were of small magnitude on an absolute scale. Collectively, these data show that anti-PD-1 and CD40 agonist antibody treatment was successful at improving the quality of the immune response in cold RFP and mixed tumor regions—contrary to our expectations that we would only observe such improvements in the hot YFP regions. However, despite the therapy-induced boost, the quality of the immune response in RFP regions remained inferior to that in YFP regions, and the absolute number of tumor-infiltrating CD8 T cells remained low. Thus, while the therapy-induced improvement in anti-tumor Th1 CD4 T cell response in RFP and mixed regions likely contributed to the blunted growth of treated tumors—which was also seen in this 6-day cohort (**Fig. 8G**)—it was largely insufficient to mediate tumor rejection.

**Figure 8.**
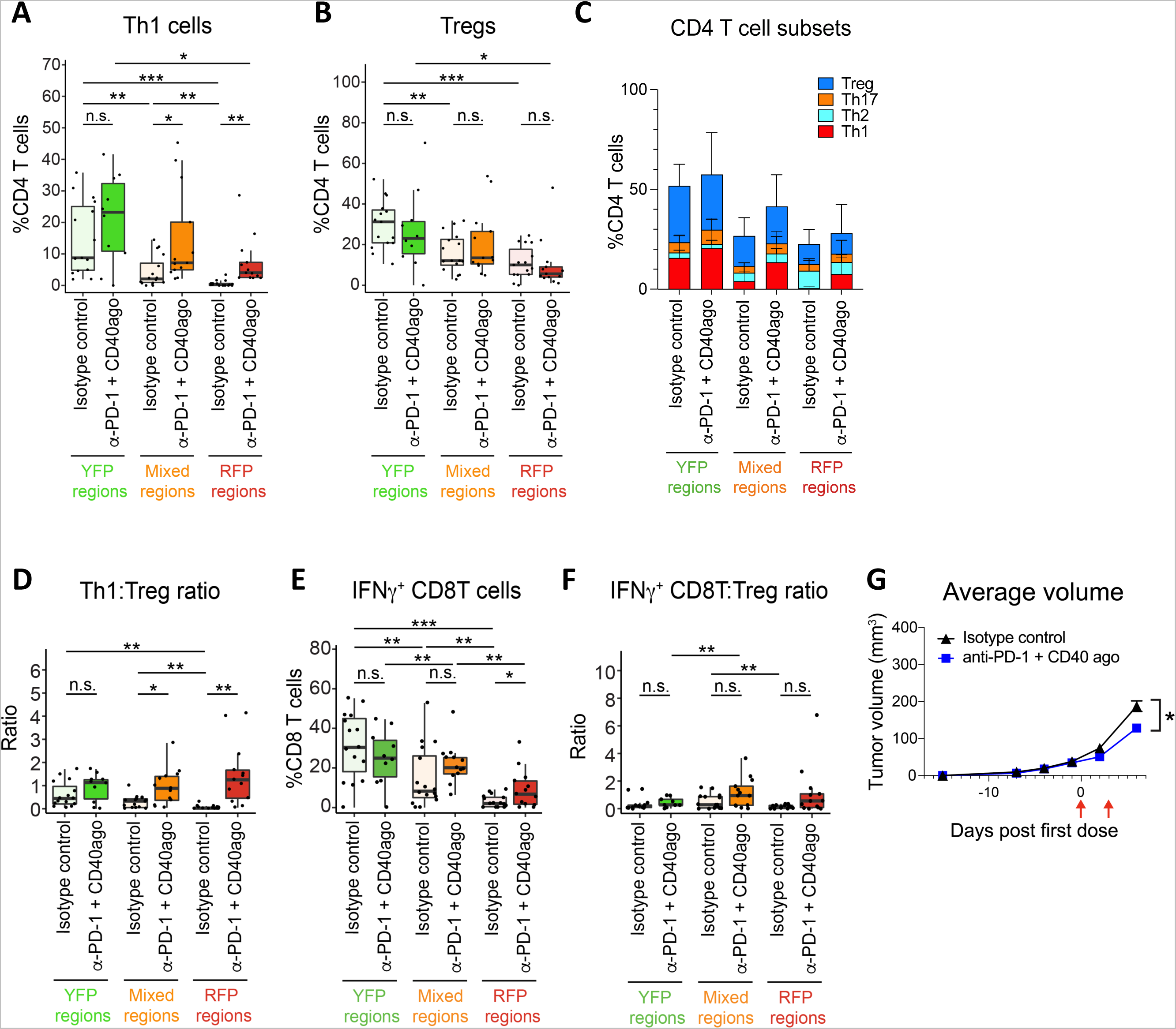
Spatial immune infiltration patterns persist in mixed-population tumors following anti-PD-1 blockade and CD40 agonist combination treatment. **(A–F)** Tumors derived from a 1:1 mixture of CIT6-YFP and CIT9-RFP cell lines were treated with α-PD-1 +CD40 agonist combination treatment or isotype control antibodies at day 0 and 3 and analyzed at day 6 for spatial T cell infiltration profile via microdissection and flow-based analysis. **(A)** Th1 cell frequency in each region of isotype control-treated and combination therapy-treated tumors, **(B)** Treg frequency, **(C)** Frequency of Th1, Th2, Th17 and Treg cells, **(D)** Th1-to-Treg ratio, **(E)** Frequency of IFN*γ*^+^ CD8 T cells and **(F)** IFN*γ*^+^ CD8 T cell to Treg ratio. **(G)** Average tumor volume of each treatment arm. Red arrows under the x axis shows the timing of treatments. Experiment was performed with at least 10 mice per treatment/tumor arm.

## Discussion

Combining fluorescently tagged pro-inflammatory (hot) and immunosuppressive (cold) squamous cell skin carcinoma tumor populations, we have developed a novel model system to interrogate the spatial heterogeneity of tumor cells and intratumoral immune cells. When combined in a 1:1 ratio, we see that an immunosuppressive tumor population has a “dominant negative” suppressive effect, in line with a previous report in a model of pancreatic cancer^15^. The overall immune phenotype of mixed-population tumors most closely resembles that of cold, immunosuppressive tumors, and exhibits a poor infiltration of CD8 T cells—which are critical to anti-tumor immunity—in all regions. Moreover, the dominant effect of the immunosuppressive tumor population is also seen locally in “mixed” regions where the two tumor populations co-exist.

However, when we look at regions locally dominated by hot tumor cells (YFP^+^ regions in our tumors), we find CD4 T cells accumulate specifically in these regions, as do inflammatory monocytes and neutrophils. Both CD4 and CD8 T cells also show better effector phenotypes in the vicinity of hot tumor cells. By contrast, immunosuppressive macrophages preferentially accumulated in the neighborhoods of cold tumor cells. Tumor cells themselves thus set up a blueprint for the spatial architecture of infiltrating immune cells, with different tumor cell populations driving distinct “micro-microenvironments” in their vicinities. Our study is the first, to our knowledge, to demonstrate a clear, reproducible link between spatial organization of infiltrating immune cells and the underlying spatial organization of tumor cells.

Tumor cell-driven spatial differences in the intratumoral immune response exist at baseline in the absence of therapy, but also persist during immunotherapy and exert an influence on the response to therapy. Regional analysis of response to anti-PD-1 blockade + anti-CD40 agonist combination immunotherapy showed a more nuanced pattern than we expected: Rather than seeing a simplistic “good” response in hot regions and “poor” response in cold regions, the relative influx of Th1 cells and IFN*γ*^+^ CD8 T cells was strongest in RFP^+^ cold regions. Nonetheless, cold regions remained more poorly infiltrated than hot regions at an absolute level, even after the therapy-induced boost, indicating that therapy did not entirely overcome the spatial patterning that existed pre-treatment. While the anti-PD-1 plus CD40 agonist combination we employed failed to cure our mixed tumors, our model system facilitated detailed region-by-region analysis of the effects and efficacy of therapy. This system thus establishes a platform on which additional drug combinations or dose regimens can be probed to iterate on how best to induce a productive tumor-wide anti-tumor immune response across all tumor regions.

We identify CX3CL1 as a mediator of a myeloid-cell-centric immunosuppressive microenvironment, and a candidate driver of the “pockets of coldness” we observe in our heterogeneous tumors. We show that CX3CL1-expressing tumor cells drive enrichment of immunosuppressive CD206^Hi^ macrophages, and depletion of inflammatory monocytes and neutrophils. In a pan-cancer analysis of human patients, high levels of CX3CL1 are associated with three specific immune cold TME archetypes, all of which show macrophage-enriched myeloid compartments and 2 of the 3 show a specific enrichment of CD206^Hi^ macrophages. The CX3CR1:CX3CL1 axis has also been found to be linked to macrophage accumulation in both skin SCC and breast tumors on a whole-tumor level^29, 30^. Further, our findings that CX3CL1 is a candidate mediator of local immunosuppressive myeloid cell organization are in line with previous reports of chemokine-driven spatial organization of immune cells within tumors^31, 32^. Of note, CX3CL1 has also been described as an “anti-tumor” chemokine, particularly in lung cancer, due to its role interacting with CX3CR1+ T cells,^26, 33–35^ highlighting that its role in tumor immunity is context-dependent and not yet fully understood. Therefore, careful investigation into CX3CL1 is essential to assess its potential therapeutic application. Of course, CX3CL1 does not fully define the immune microenvironment. Future work, in addition to more deeply investigating the role of CX3CL1, is also expected to uncover additional pathways by which tumor cells shape their local micro-microenvironments and influence responses to immunotherapy on a highly local spatial scale.

## Materials and methods

### Carcinogenesis and cell line generation

To induce tumors, *K5-CreER^T2^-Confetti FVB/N* mice were treated with 25 mg DMBA followed by TPA (200 µl of a 10^−4^ M solution in acetone) two times a week for 20 weeks, as described^19^. To generate cell lines, carcinomas were resected at a size of >1 cm in longest diameter, finely chopped, digested in DMEM containing Collagenase I and IV and DNase I for 45 minutes at 37 °C, washed with PBS, plated in supplemented DMEM (high-glucose DMEM (Gibco, catalog# 11995065) plus 10% heat-inactivated FBS (Gibco), 2mM L-glutamine, with 100 units/ml penicillin, 100 µg/ml streptomycin, and 2.5 µg/ml amphotericin B (Gibco)), and passaged until cells stably grow in culture. The established cell lines were cultured in the supplemented DMEM for all experiments.

### Cell line labeling

Unlabeled cell lines were treated with adenoviral Cre recombinase (Vector Biolabs catalog #1045) to induce labeling with Confetti fluorescent proteins. YFP- and RFP-expressing cells were sorted via fluorescence-activated cell sorter and further grown in vitro to establish CIT6-YFP and CIT9-RFP cell lines.

### In vivo experiments

A total of 1.25 × 10^5^ cells were injected subcutaneously into the dorsal flank of 8 to 16 week-old male and female *Confetti*-homozygous *FVB/N* mice. Approximately 21-28 days later, when tumors measured ∼1cm in diameter, tumors were dissected and digested as described above, and subsequently used in flow cytometric analysis. For treatment experiments, when tumors measured approximately 5mm in diameter, mice were injected intraperitoneally with one dose (CD40 agonistic antibody) or two doses (anti-PD-1 antibody) of therapeutic antibodies, administered at a three day interval. Antibodies used were rat anti-mouse PD-1 IgG2a antibody (RMP1-14, BioXCell), rat anti-CD40 agonist IgG2a antibody (FGK4.5, BioXCell), and rat IgG2a isotype control (2A3; BioXCell) at 200μg per antibody per dose. Tumors were resected at 1cm in diameter and used for downstream analyses, except in 6-day post-treatment analyses where tumors were resected 6 days after treatment initiation. All animal experiments were approved by the University of California San Francisco Laboratory Animal Resource Center (IUCAC; AN187679). This work complies with all the relevant ethical regulations regarding animal research.

### Flow cytometric immune profiling and macrophage cell sorting

For whole tumor analysis and sorting, 5 million cells were stained with antibodies. For spatial immune profiling, tumors were embedded in 2% agarose gel, sliced into 400µm section by a Leica Vibratome, microdissected into YFP, RFP and mixed regions using surgical scalpel while visualizing each region on MVX10 fluorescent stereoscope. Multiple pieces of each region from a single tumor were pooled for digestion, followed by flow cytometric analysis. Cell sorting was performed on FACS Aria III (BD Biosciences). Antibodies used for flow cytometric analyses and sorting are listed in Table 2.

### Immunofluorescence imaging

After half of the tumors were processed for flow cytometric analysis, the remaining halves were fixed in 4% paraformaldehyde for 2 hours, dehydrated in PBS with 30% sucrose overnight at 4C°, embedded in Tissue-tek O.C.T. compound (Sakura Finetek USA) and stored at −80C°. Embedded blocks were cryosectioned into 8µm sections using Leica CM3050 S Cryostat (Leica Biosystems) and stained with rabbit anti-mouse CD3 antibody (ab5690, Abcam) overnight, followed by stain with secondary anti-rabbit Alexa Fluor 647 antibody (Invitrogen) for 1 hour at room temperature and DAPI stain for 5 minutes. Fluorescence image was taken on microscope (DMi8, Leica Biosystems) using Leica Application Suite X (Leica Biosystems) imaging software, and fraction of YFP+ and RFP+ cells as well as the number of CD3+ cells in each tile were quantified using CellProfiler ver. 3.1.9.

### Overexpression of cytokines and chemokines

Mouse *Cx3cl1* gene was amplified using forward primer 5’-aactcgagatggctccctcgccgctcg-3’ and reverse primer 5’-ttccgcggtcacactggcaccaggacgta-3’, cDNAs generated from CIT9 cell line and Q5® High-Fidelity DNA polymerase (NEB). MSCV 2.2-IRES-CFP retroviral vector was generated by replacing GFP of MSCV 2.2-IRES-GFP vector (obtained from James Carlyle lab at University of Toronto) with CFP. Using restriction enzymes XhoI and SacII, *Cx3cl1* coding sequencing was cloned into MSCV 2.2-IRES-GFP vector. Proviral vectors Gag/Pol and VSV-G were co-transfected with Cx3cl1-IRES-GFP MSCV2.2 vector using lipofectamine 3000 (Invitrogen) into HEK293T cells. 48 hours later, the viruses were collected, mixed with CIT6-YFP cells and spinfected at 800g for 90 minutes. YFP^+^ CFP^+^ transduced cells were sorted 72 hours post-transduction.

### ZipSeq spatial transcriptomics analysis

Mixed-population tumors were harvested at 1cm in diameter and sliced into 200µm section in 2% agarose using vibratome. Section was incubated with CD45 antibody linked to Cy5-conjugated photocaged oligo for 1 hour at 4C°, followed by patterned illumination of a user-defined region of interest^25^ and incubation with a unique ZipCode. The process was repeated for YFP, RFP and mixed regions. The section then was digested in a Collagenase I and IV blend for 30 minutes at room temperature, and sorted for live Cy5^+^ cells using FACS Aria II. washed in PBS + 0.04% BSA and then encapsulated following 10× Genomics specifications for 3’ v3 chemistry with target cell number of 8,000.

Libraries were sequenced with a target number of 30,000 reads per cell for gene expression and 3,000 reads per cell for ZipCode barcodes on a Illumina NovaSeq. Demultiplexed fastqs were aligned using CellRanger to the mm10 Ensembl 93 reference genome. Count matrices were used to generate Seurat objects in R keeping only cells with at least 200 detected genes and genes found in at least 3 cells. Cells were further filtered based on a minimum 500 gene cutoff and maximum 15% mitochondrial reads. ZipCode counts were normalized using centered log ratio transform and then assigned a dominant ZipCode identity (as in ^25^). Cells were then passed through SCTransform regressing for percent mitochondrial reads. Following PCA analysis on the top variable genes, the first 18 principal components were used as inputs for UMAP dimensional reduction and clustering via the FindNeighbors and FindClusters commands. For pathway analysis, lists of genes with average Log_2_ fold change >1 for each cluster were used as input for GO Biological Process 2021 repository on EnrichR (https://maayanlab.cloud/Enrichr/).

### Cytokine array

70-150mg of tumor pieces were chopped into small pieces in RIPA buffer containing proteinase inhibitor (Cell BioLabs) and homogenized using OctoMACS (Miltenyi Biotec). The homogenate was centrifuged at 400g for 5 minutes followed by further centrifugation at 10,000g for 10 minutes at 4C°. The supernatant was collected and stored at −80C°. Protein concentration was measured using DC protein assay kit (BioRad). The sample concentrations were normalized prior to the submission for the analysis by Eve Technologies DM-44 mouse Discovery Assay panel.

### *In vitro* T cell suppression co-culture assay

CD8 T cells were isolated from the lymph node of naïve mice with unactivated Confetti cassette using EasySep™ Mouse CD8^+^ T Cell Isolation Kit (StemCell Technologies) and macrophages were sorted from the tumors via FACS sorting. CD8 T cells were labeled with CFSE and co-cultured with the macrophages at varying ratios in the presence of Mouse T-Activator CD3/CD28 Dynabeads™ (Gibco) and analyzed for division and T cell activation markers.

### Immune cell abundance analysis for human RNAseq dataset

For analysis of human tumors, T cell, macrophage and monocyte scores were based on previously calculated scores using an average expression level of genes that are uniquely expressed in human T cells, macrophages and monocytes^28^. The CD206^Hi^ macrophage score was calculated by identifying a combination of four genes that are uniquely expressed in the CD206^Hi^ macrophage cluster (cluster 3) of our ZipSeq dataset (*Hmox1, Cx3cr1, Folr2 and F13a1*). This score was normalized against the average expression level of four genes (*Ccr2, Mgl2, Nr4a2 and Ciita*), which were expressed in other macrophages but were not expressed in the CD206^Hi^ macrophage cluster.

### Statistical analysis

Data were analyzed using Prism 9 (GraphPad) with either Student’s t-test for experiments with two arms, one-way ANOVA analysis for experiments with one variable and more than 3 experiment conditions, or two-way ANOVA for experiments with two variables and more than 3 experiment conditions. Statistical tests for spatial analyses (Fig. 3 and 8) were done using Student’s t-test. For Kaplan-Meier survival analyses, log-rank test was used. Graphs show mean ± SEM or mean ± SD; *, *p* < 0.05; **, *p* < 0.01; ***, *p* < 0.001.

## Acknowledgements

This work was supported by the UCSF Program for Breakthrough Biomedical Research (PBBR) Sandler Fellowship, US National Cancer Institute (NCI) grant R21CA264599, the Huntsman Cancer Institute Cancer Center Support Grant P30CA040214, the Parker Institute for Cancer Immunotherapy, the American Cancer Society, and The Cancer League. We thank the UCSF Biological Imaging Development CoLab and UCSF Parnassus Flow CoLab, RRID:SCR_018206.

## Author Contributions

M.T., L.L., C.L.-S., and M.Q.R. designed the project and experiments. M.T., L.L. and C.L.-S. led all experiments and data analyses. K.H. and M.T. carried out the ZipSeq single cell transcriptomics experiments and analysis, with support from M.K. A.J.C. and B.S. conducted the analysis of human RNAseq and immune archetype datasets. M.T. and M.Q.R. wrote the manuscript, with input from other co-authors. K.K., D.S., K.N., and Z.A. assisted with experiments. L.F. provided guidance to the studies.

