## Supplementary Figures for "Tumor cell heterogeneity drives spatial organization of the intratumoral immune response in squamous cell skin carcinoma"

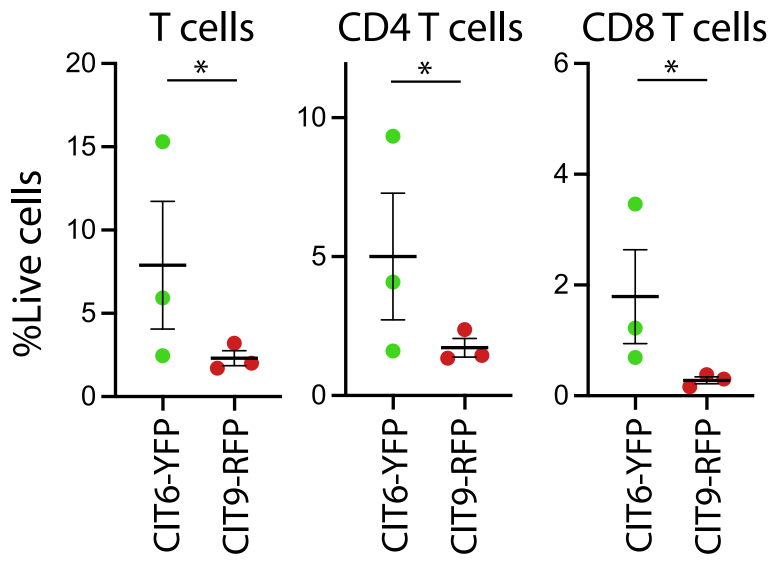


**Supp. Fig. 1. Abundance of total, CD4 and CD8 T cells as fraction of live cells in CIT6-YFP and CIT9-RFP subcutaneous tumors.**

CIT6-YFP and CIT9-RFP tumors were injected into homozygous *Confetti* mice and harvested at 10mm for T cell analysis by flow cytometry.

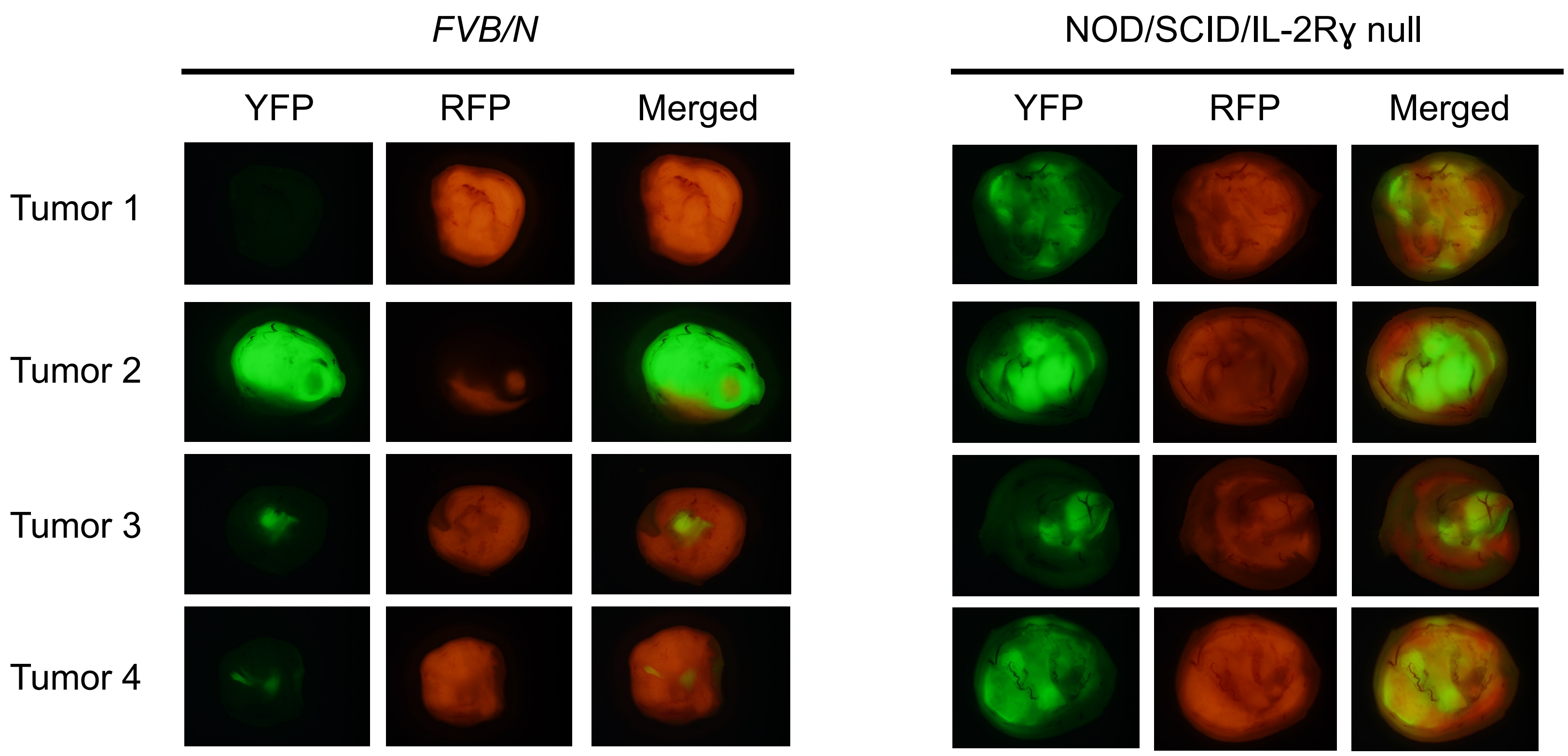


**Supp. Fig. 2 Comparison of YFP and RFP signals in mixed-population tumors between FVB/N and NSG mice.** Fluorescent stereomicroscope whole-tumor images of mixed-population tumors formed in *FVB/N* mice and NOD/SCID/IL-2Rɣ null mice.

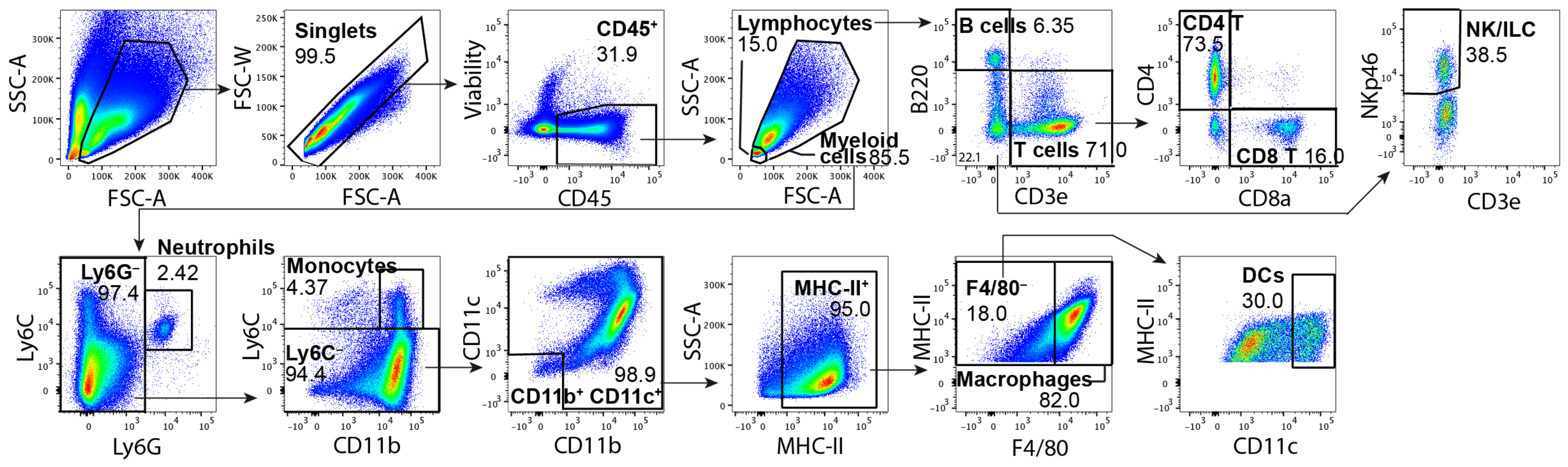


**Supp. Fig. 3. Flow cytometric gating strategy of main immune cell types.**
Immune analysis of subcutaneous CIT6-YFP tumor is shown as a representative data.

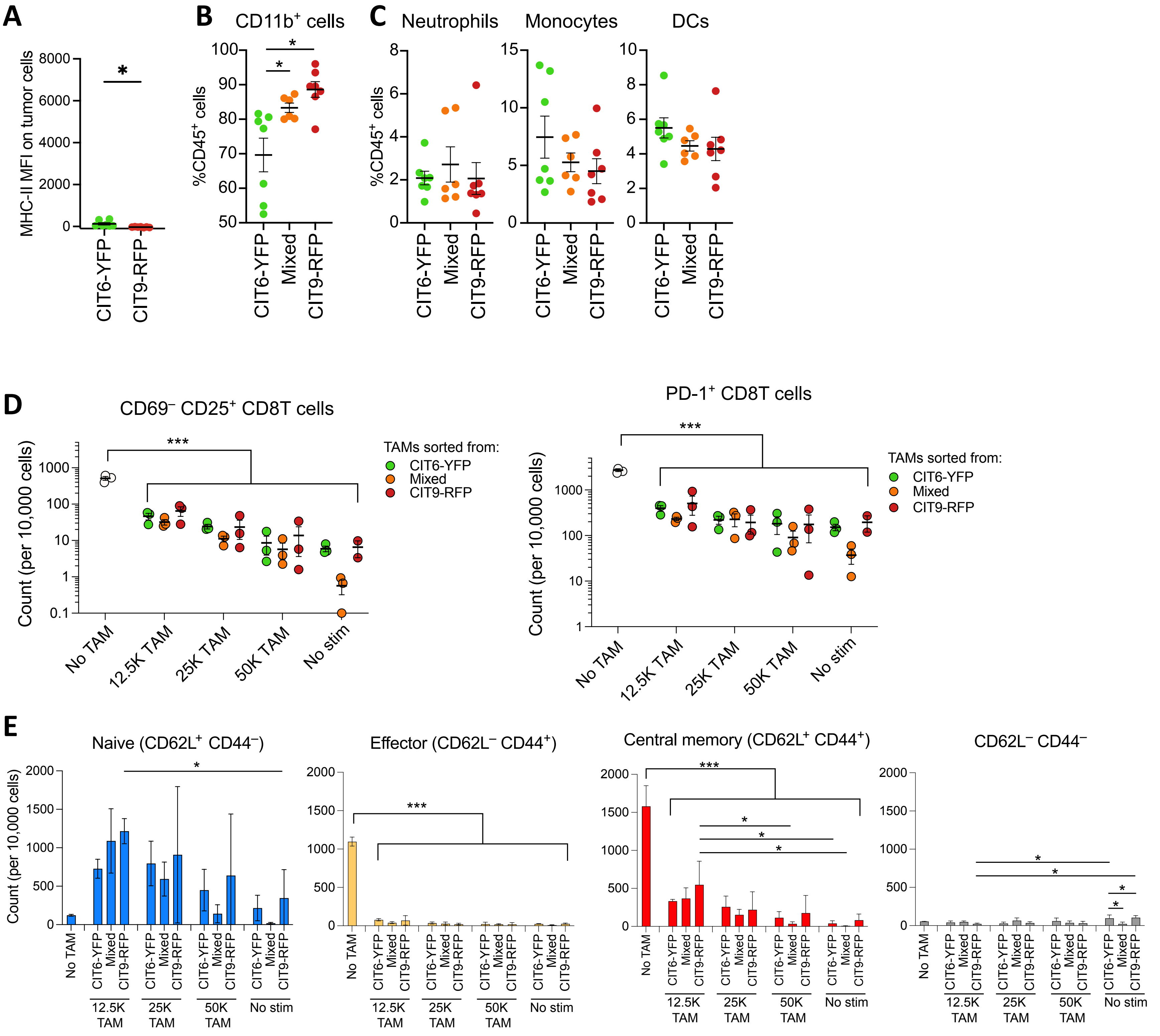


**Supp. Fig. 4 Myeloid cell infiltration and macrophage function in CIT6-YFP, CIT9-RFP and mixed-population tumors.**
**(A)** Expression of MHC-II (I-A/I-E) on tumor cells in CIT6-YFP and CIT9-RFP tumors. **(B)** Frequency of CD11b$^{+}$ cells. **(C)** Frequency of neutrophils, monocytes and DCs. **(D)** Frequency of CD69$^{-}$ CD25$^{+}$ and PD-1$^{+}$ activated T cells in *in vitro* co-culture assay of CD8 T cells and macrophages sorted from CIT6-YFP, CIT9-RFP and mixed-population tumors. Data are a representation of 2 independent experiments. **(E)** Frequency of effector T cell subsets broken down by subset to show statistical test results.

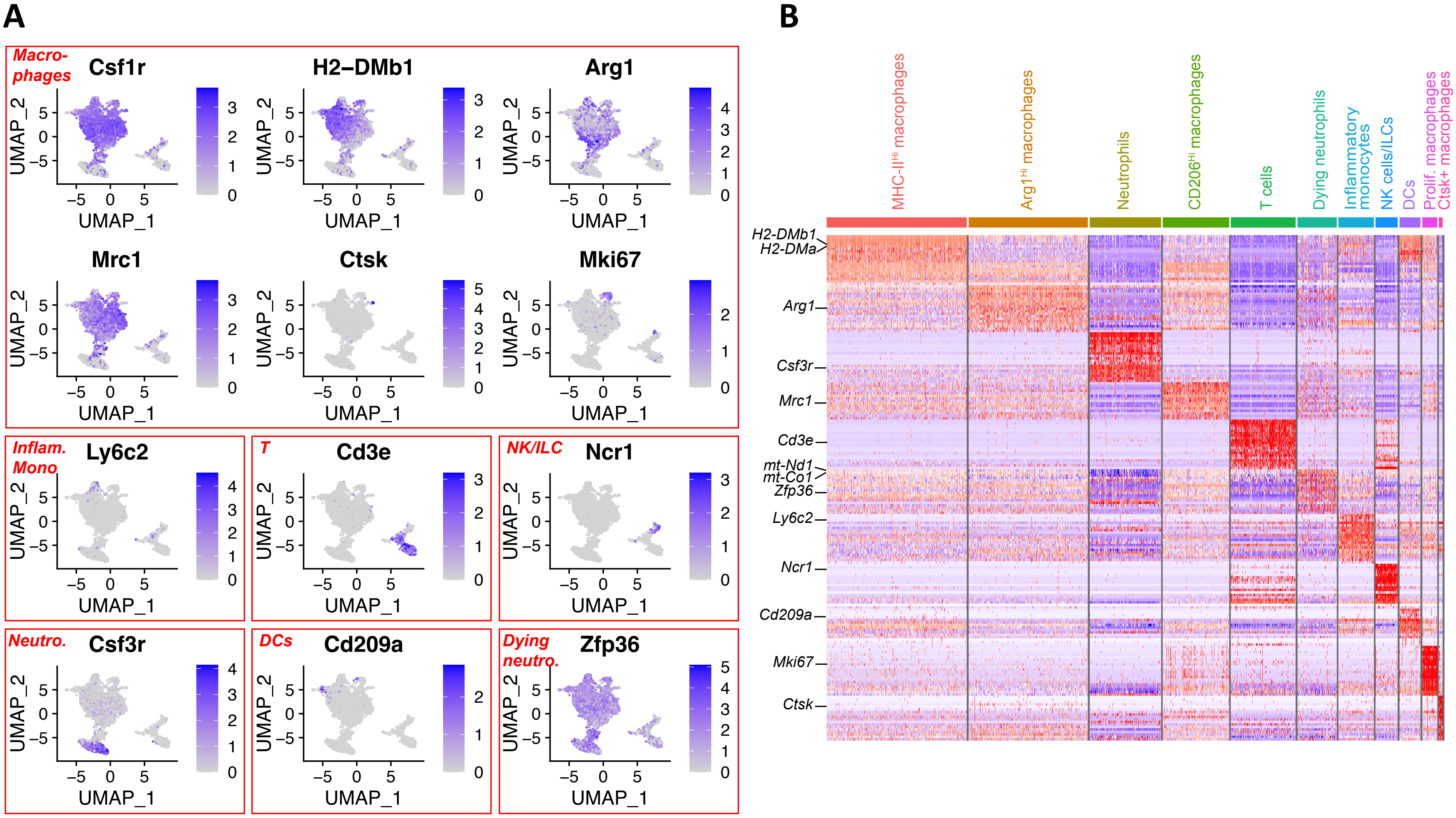


**Supp. Fig. 5. Annotation of clusters detected in ZipSeq data based on expression of well-established immune cell markers.**
**(A)** Feature plots overlaid on UMAP representation for *Csfr1*, marking macrophage clusters, and and a marker gene for each cluster of the tumor analyzed by ZipSeq in Fig. 4. **(B)** Heatmap showing relative average expression of the 20 most strongly enriched genes for each cluster versus all others.

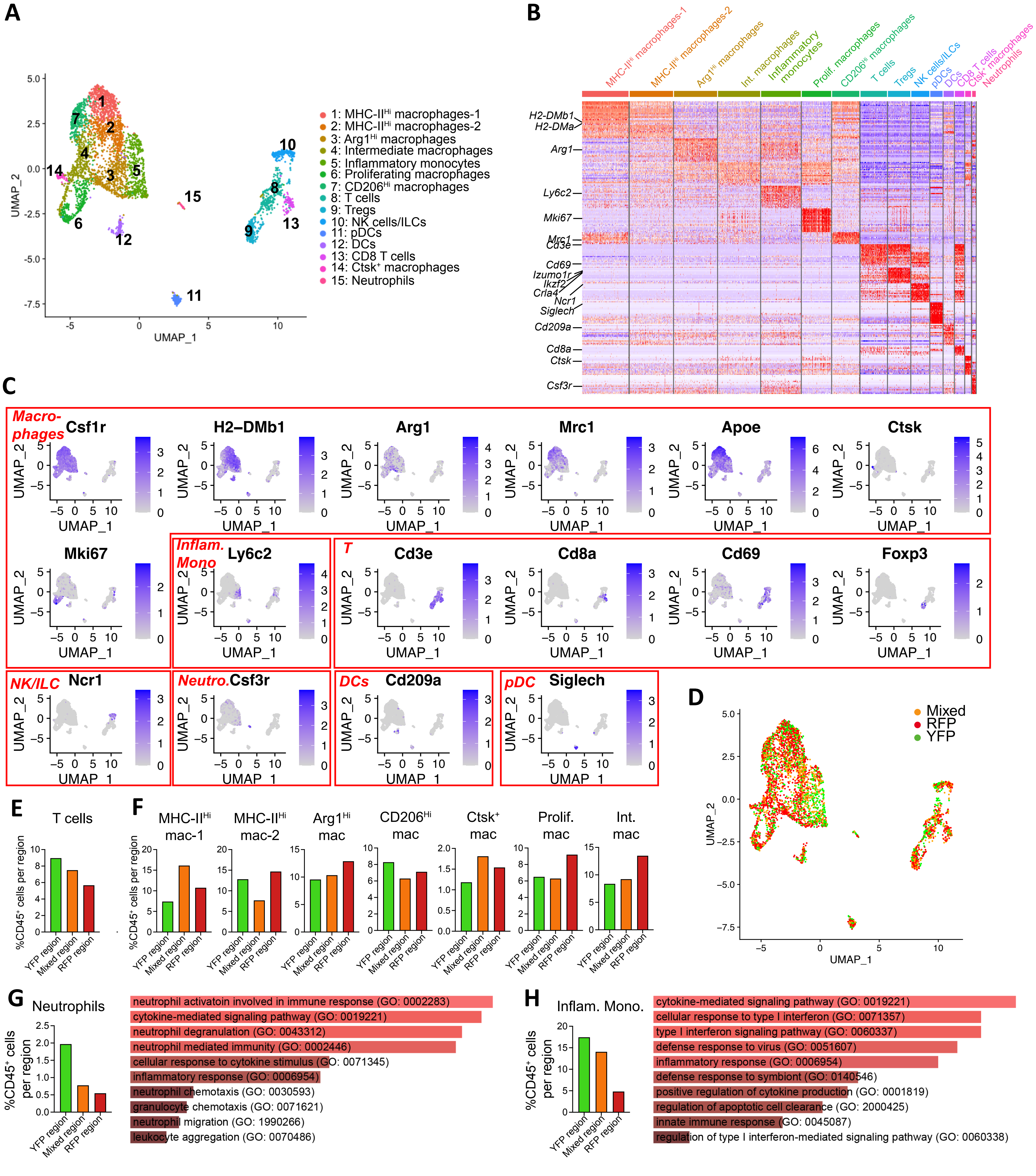


**Supp. Fig. 6. ZipSeq analysis of an additional mixed-population tumor.**
**(A)** UMAP representation of zipcode-labeled cells with cluster overlay. n = 4,186 cells, nYFP = 1,015, nRFP = 2,011, nMixed = 1,160. **(B)** Heatmap showing the 20 most enriched genes in each cluster. Marker gene(s) for each cluster are labeled on the left of the heatmap. **(C)** Feature plots overlaid on UMAP representation for a gene marking macrophage clusters (*Csf1r*) and a marker gene of each cluster. **(D)** UMAP representation of zipcode-labeled cells with zipcode identity overlaid. **(E)** Abundance of cells belonging to the T cell cluster , calculated as percentage of total immune cells in each region. **(F)** Abundance of cells belonging to seven macrophage clusters (clusters 1, 2, 3, 4, 6, 7, and 14), calculated as percentage of total immune cells in each region. **(G)** Percentage of cells belonging to the neutrophil cluster (cluster 15) in each region and pathway analysis showing gene families enriched in the neutrophil cluster. **(H)** Percentage of cells belonging to the monocyte cluster (cluster 5) and pathway analysis of genes enriched in the monocyte cluster.

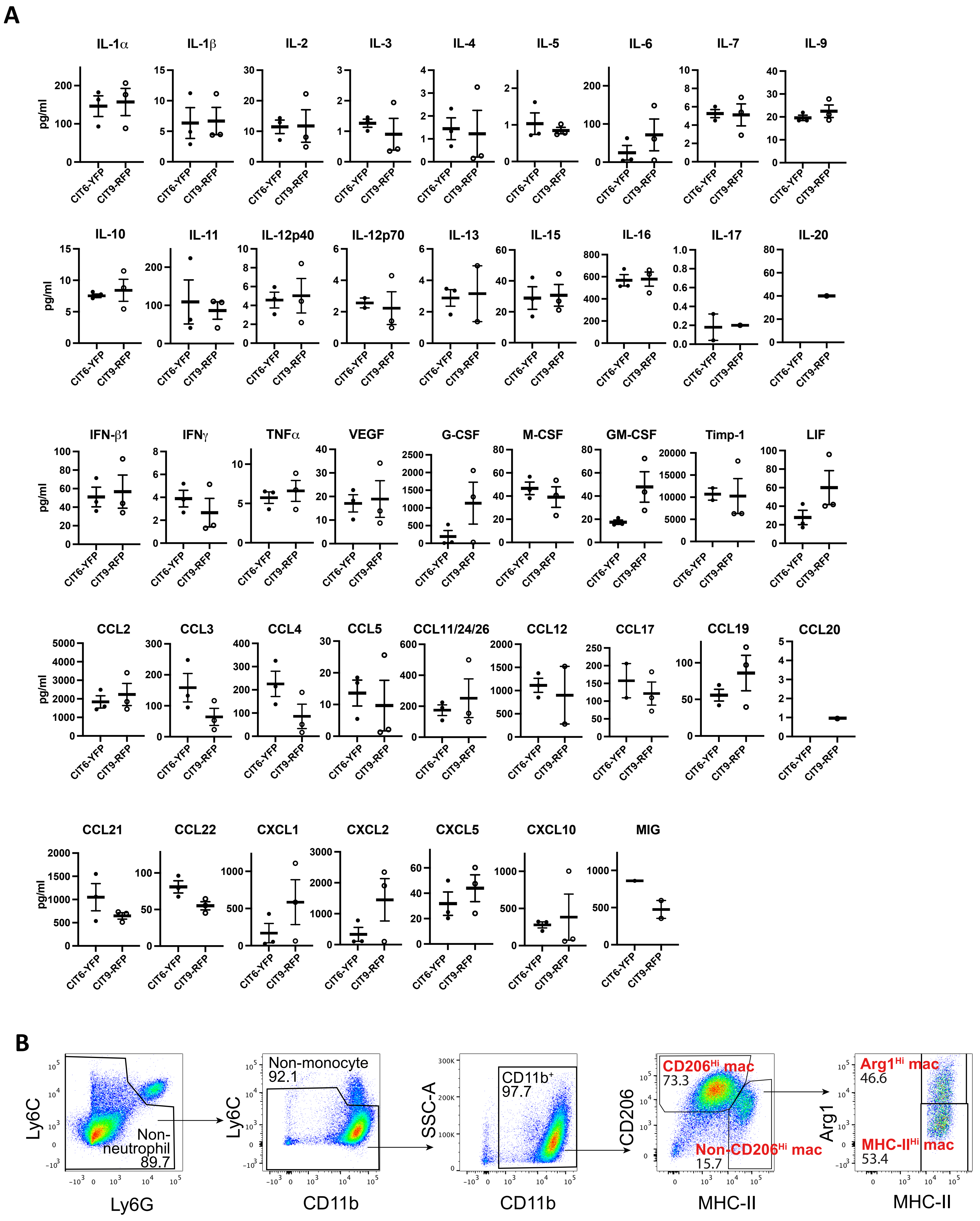
A
IL-1α
IL-1β
IL-2
IL-3
IL-4
IL-5
IL-6
IL-7
IL-9
IL-10
IL-11
IL-12p40
IL-12p70
IL-13
IL-15
IL-16
IL-17
IL-20
IFN-β1
IFNγ
TNFα
VEGF
G-CSF
M-CSF
GM-CSF
Timp-1
LIF
CCL2
CCL3
CCL4
CCL5
CCL11/24/26
CCL12
CCL17
CCL19
CCL20
CCL21
CCL22
CXCL1
CXCL2
CXCL5
CXCL10
MIG
pg/ml
CIT6-YFP
CIT9-RFP
B
Ly6C
Ly6G
Non-neutrophil
89.7
Non-monocyte
92.1
CD11b
SSC-A
CD11b+
97.7
CD206
MHC-II
CD206Hi mac
73.3
Non-CD206Hi mac
15.7
Arg1
Arg1Hi mac
46.6
MHC-IIHi mac
53.4

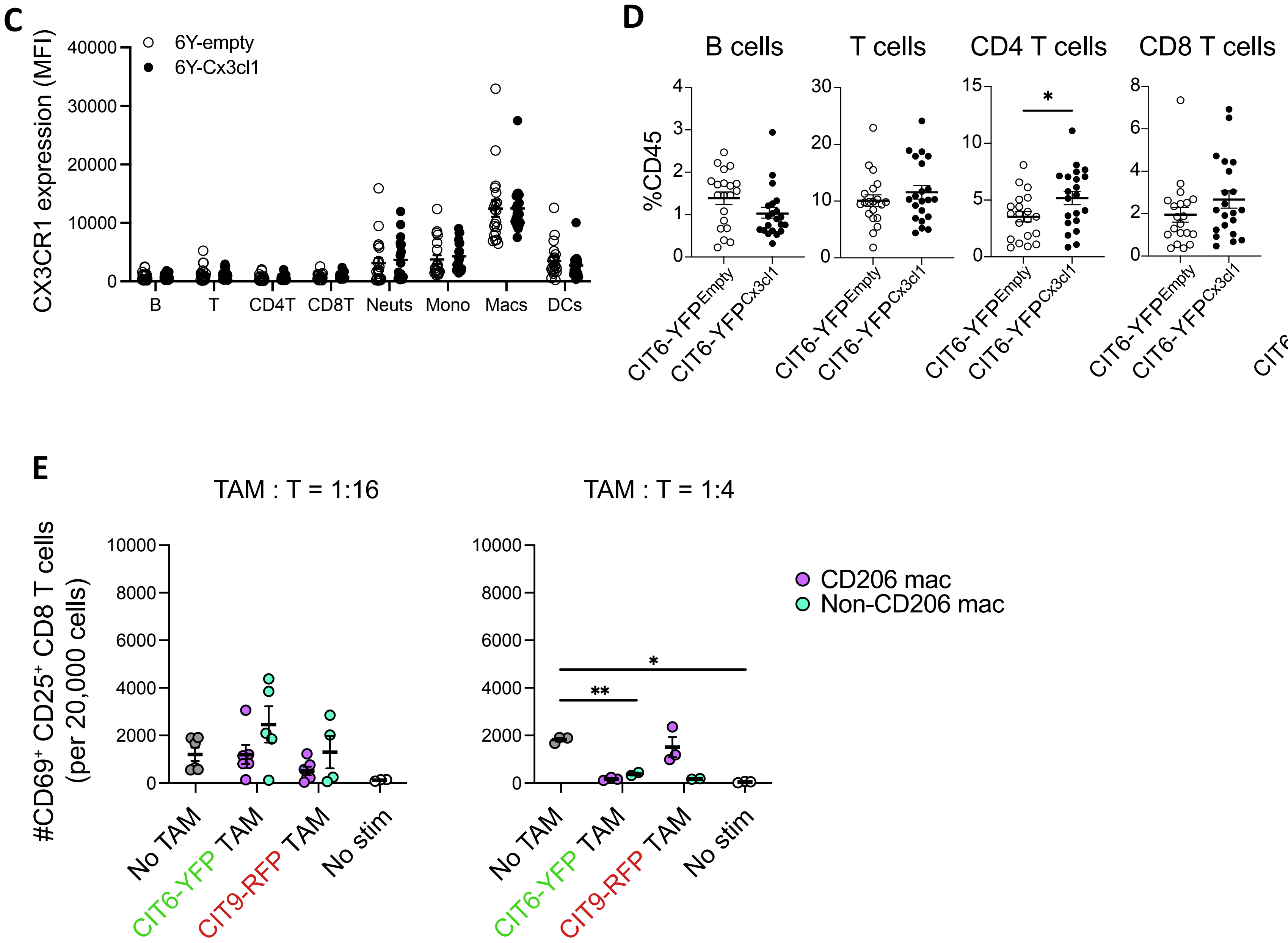


**Supp. Fig. 7 Characterization of infiltrating immune cells in Cx3cl1-overexpressing CIT6-YFP tumors.**
**(A)** Protein expression of 44 cytokine and chemokines in CIT6-YFP and CIT9-RFP tumors measured by ELISA. **(B)** Gating strategy of identifying MHC-II$^{Hi}$, Arg1$^{Hi}$ and CD206$^{Hi}$ macrophage subsets used for flow cytometric analysis and FACS sorting. CD206$^{Hi}$ TAM and non-CD206$^{Hi}$ populations on the fourth plot were used for co-culture T cell suppression assay. **(C)** CX3CR1 protein expression on immune cell types shown as MFI from flow cytometric analysis. **(D)** Frequency of lymphocytes and DCs in Cx3cl1-overexpressing and empty vector-expressing CIT6-YFP tumors. **(E)** Number of activated (CD69$^{+}$ CD25$^{+}$) CD8 T cells at TAM:T = 1:16 and 1:4 co-culture ratios.

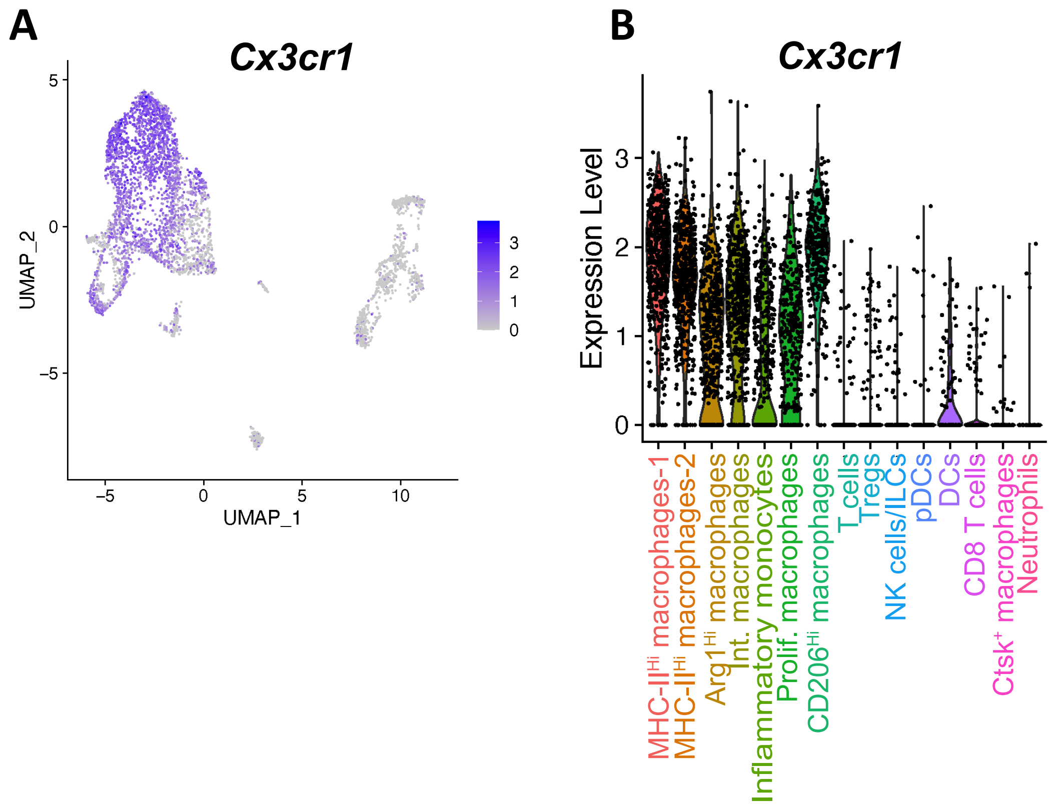


**Supp. Fig. 8 *Cx3cr1* expression distribution across clusters in a mixed-population tumor analyzed by ZipSeq in Supp.Fig. 6.**
**(A)** Feature plot overlaid on UMAP representation for *Cx3cr1* gene. **(B)** Violin plot showing *Cx3cr1* distribution based on cluster identity.

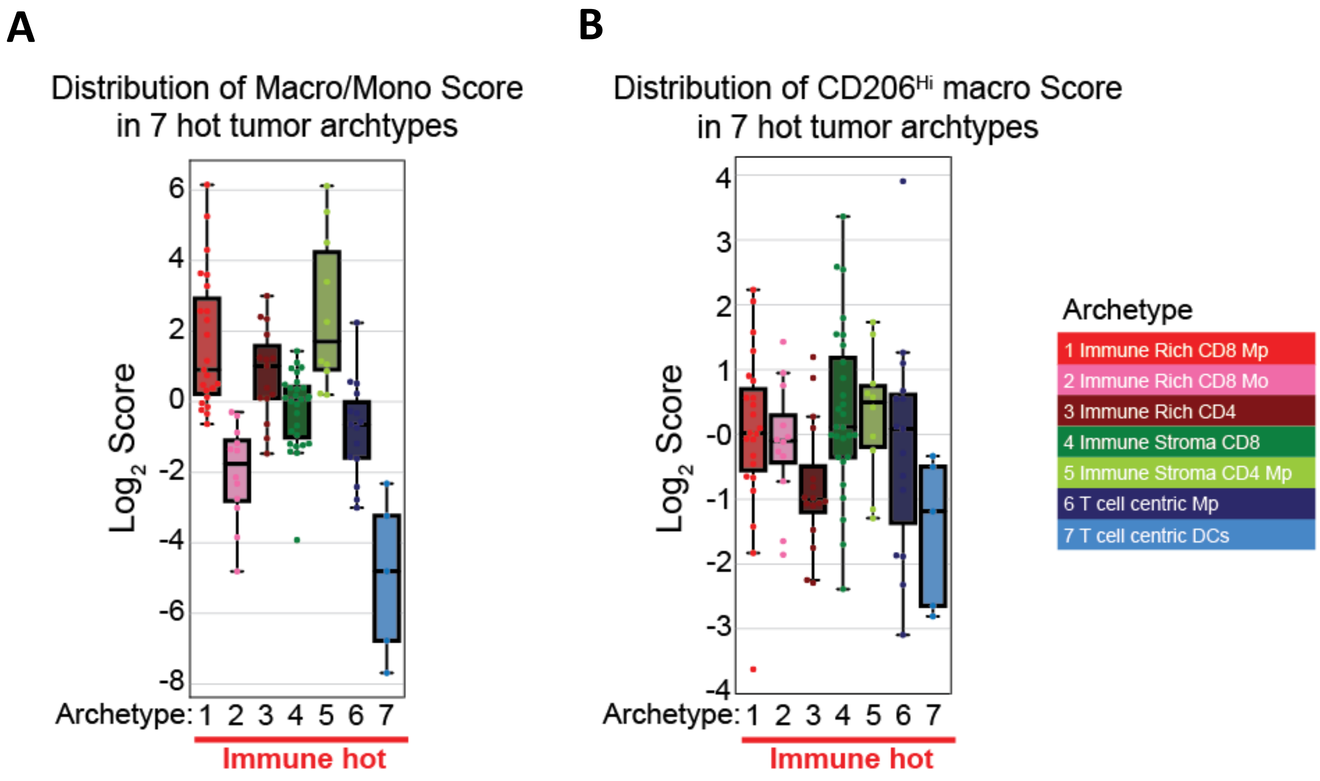


**Supp Fig. 9 Macrophage:monocyte ratio and CD206Hi macrophage scores in tumors with hot immune archetypes.**
Text text text

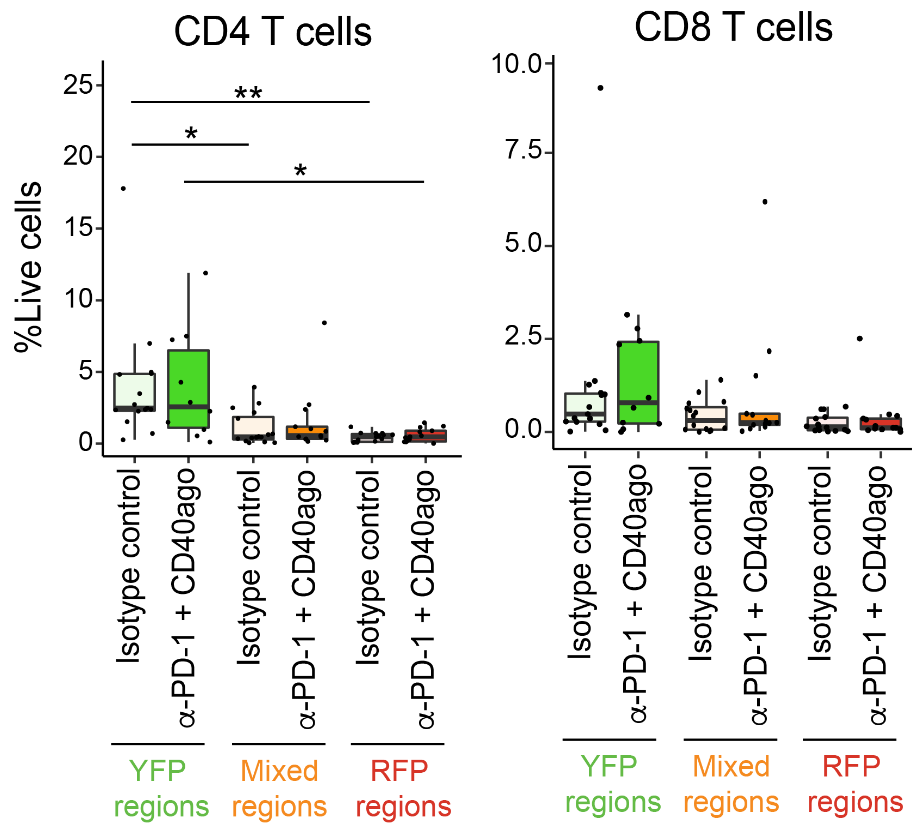


**Supp. Fig. 10 Spatial infiltration pattern of CD4 T and CD8 T cells in mixed-population tumors post anti-PD-1 blockade and CD40 agonist combination treatment.**

Total CD4 T cell ($CD45^+$ $CD3^+$ $CD4^+$) and total CD8 T cell ($CD45^+$ $CD3^+$ $CD8^+$) frequencies in each region of isotype control-treated and combination therapy-treated tumors analyzed at day 6 post treatment initiation.
