## Supplementary material for "Tumor cell heterogeneity drives spatial organization of the intratumoral immune response in squamous cell skin carcinoma": Table 2

| Marker | Fluorophore | Clone | Manufacturer |
|---|---|---|---|
| CD45 | AlexaFluor700 | 30-F11 | BioLegend |
| F4/80 | BV421 | BM8 | BioLegend |
| CD11c | BV605 | N418 | BioLegend |
| Ly6C | BV650 | HK1.4 | BioLegend |
| CD11b | BV711 | M1/70 | BioLegend |
| CD8a | BV785 | 53-6.7 | BioLegend |
| CD3e | BUV395 | 145-2C11 | BD Biosciences |
| CD4 | BUV563 | GK1.5 | BD Biosciences |
| B220 | BUV661 | RA3-6B2 | BD Biosciences |
| Ly6G | PE-CF594 | 1A8 | BD Biosciences |
| MHC II | PE-Cy7 | M5/114.15.2 | BioLegend |
| CX3CR1 | APC | SA011F11 | BioLegend |
| CX3CR1 | PE | SA011F11 | BioLegend |
| F4/80 | APC | BM8 | BioLegend |
| Arg1 | eFluor450 | A1exF5 | Invitrogen |
| CD206 | BV785 | C068C2 | BioLegend |
| CD206 | PE | C068C2 | BioLegend |
| TNF$\alpha$ | PerCPCy5.5 | MP6-XT22 | BioLegend |
| Foxp3 | APC | FJK-16s | Invitrogen |
| CD25 | BV421 | PC61 | BioLegend |
| PD-1 | BV605 | 29F.1A12 | BioLegend |
| IL17A | BV650 | TC11-18H10.1 | BioLegend |
| IL-4 | BV786 | 11B11 | BD Biosciences |
| IFN$\gamma$ | BUV737 | XMG1.2 | BioLegend |
| CD39 | PE-Dazzle594 | Duha59 | BioLegend |
| CD69 | BV605 | H1.2F3 | BioLegend |
| CD44 | BV785 | IM7 | BioLegend |
| CD62L | BUV737 | MEL-14 | BD Biosciences |
| NKp46 | BV421 | 29A1.4 | BioLegend |
| MHC-I (H-2Kq) | BV421 | KH11 | BD Biosciences |
| MHC-I (H-2Dq, Lq) | Biotin | KH117 | BD Biosciences |
| CD16/32 | N/A | 2.4G2 | Tonbo Biosciences |
| Live/dead | Fixable near-IR | N/A | Invitrogen |
